# Quantifying microbially mediated fitness differences reveals the tendency for plant-soil feedbacks to drive species exclusion among California annual plants

**DOI:** 10.1101/2020.02.13.948679

**Authors:** Gaurav S. Kandlikar, Xinyi Yan, Jonathan M. Levine, Nathan J.B. Kraft

**Author notes:** ^*^Author for correspondence: Gaurav Kandlikar. Coauthor contact information: Xinyi Yan, Jonathan M. Levine, Nathan J.B. Kraft.

## Abstract

Soil microorganisms influence a variety of processes in plant communities. Many theoretical and empirical studies have shown that dynamic feedbacks between plants and soil microbes can stabilize plant coexistence by generating negative frequency-dependent plant population dynamics. However, inferring the net effects of soil microbes on plant coexistence requires also quantifying the degree to which they provide one species an average fitness advantage, an effect that has received little empirical attention. We conducted a greenhouse study to quantify microbially mediated stabilization and fitness differences among fifteen pairs of annual plants that co-occur in southern California grasslands. We found that although soil microbes frequently generate negative frequency-dependent dynamics that stabilize plant interactions, they simultaneously mediate large average fitness differences between species. The net result is that if the plant species are otherwise competitively equivalent, the impact of plant-soil feedbacks is often to favor species exclusion over coexistence, a result that only becomes evident by quantifying the microbially mediated fitness difference. Our work highlights that comparing the stabilizing effects of plant-soil feedbacks to the fitness difference they generate is essential for understanding the influence of soil microbes on plant diversity.

## Introduction

The dynamics of plants and soil microorganisms are tightly intertwined. The composition of soil microbial communities responds strongly to different plant species, in large part due to interspecific differences in root architecture and exudate profile (Berg and Smalla 2009). Soil microorganisms in turn influence the growth of plant species, with the direction and magnitude of their effect determined both by the composition of the microbial community and by the genetic and functional characteristics of the plants (Laliberté et al. 2014, Keller and Lau 2018). These plant-soil feedbacks can have important consequences for various processes in plant communities (van der Putten et al. 2016), including species coexistence.

The effects of soil microorganisms on plant species coexistence are typically studied in the context of a theoretical framework developed by Bever et al. (1997). This framework isolated the effects of soil microbes by modeling the dynamics of plant species that differ only in the soil microbial communities they cultivate and in how their growth is influenced by these cultivated communities. Bever et al. (1997) showed that the soil microbial community stabilizes plant interactions when the microbial community cultivated by each plant species limits the growth of the cultivating plant species more (or benefits it less) than that of the other plant species in the system. When this condition is satisfied, the microbial community generates a relative advantage in favor of species that fall in abundance and limits the abundant species, thereby generating negative frequency-dependent plant population dynamics that should promote species coexistence. Alternatively, plant-soil feedbacks can destabilize plant interactions if they create positive feedback loops that favor abundant species over species that fall in abundance. Results from numerous empirical studies motivated by this framework indicate that plant-soil feedbacks often drive such frequency-dependent plant population dynamics. Feedbacks generally stabilize interactions among plant species that associate with similar mycorrhizal guilds, are distantly related, or are interacting in both species’ native range (reviewed in Crawford et al. (2019)).

Although plant-soil feedback research has emphasized the potential for these interactions to stabilize plant interactions, we still lack a general understanding of whether soil microbes generally favor plant coexistence or species exclusion. This is in part because inferring the coexistence consequences of plant-soil feedbacks from their (de)stabilizing effects alone, and not comparing this effect to the degree to which soil microbes mediate an average fitness difference between plant species, can lead to false conclusions about how soil microbes influence plant diversity (Chesson 2000, Kandlikar et al. 2019). This microbially mediated fitness difference reflects plant species’ variation in their average sensitivity to the pathogenic or mutualistic soil microbes cultivated by both conspecifics and heterospecifics. Importantly, quantifying the degree to which plant-soil feedbacks drive average fitness differences between species requires measuring plant growth with a reference uncultivated soil microbial community (Kandlikar et al. 2019), an experimental treatment that is excluded in most studies of plant-soil feedback.

To fully evaluate the influence of microbially mediated stabilization and fitness differences on plant coexistence, we conducted a two-phased experiment (Bever et al. 2010) in which we first grew monocultures of six plant species to cultivate their characteristic soil microbial community, and then measured the growth of each species in soils inoculated with a distinct microbial community – including a field-collected microbial inoculum that was not cultivated by any of the focal species. We used these data to estimate the key parameters from Bever et al. (1997)’s model of plant-soil feedback. Then, we quantified the degree to which soil microbes stabilize or destabilize pairwise plant interactions and the degree to which they drive average fitness differences using metrics derived in Kandlikar et al. (2019). Our study shows that even when plant-soil feedbacks stabilize species interactions, their net effect can be to favor species exclusion if they simultaneously drive strong fitness differences between plant species.

## Methods

### Study system

We studied the effects of plant-soil feedbacks on the pairwise interactions of six annual plant species: *Acmispon wrangelianus* (Fabaceae), *Festuca microstachys* (Poaceae), *Hordeum murinum* (Poaceae), *Plantago erecta* (Plantaginaceae), *Salvia columbariae* (Lamiaceae), and *Uropappus lindleyi* (Asteraceae). These species co-occur in the winter annual plant community in the University of California Sedgwick Reserve in Santa Barbara County, California, USA (34 41’ N, 120 02’ W, 290-790m above sea level). This region experiences a Mediterranean climate of cool, wet winters (October-May mean temperature = 13.5°C, mean monthly precipitation = 164mm) and hot, dry summers (June-September mean temperature = 20°C, mean monthly precipitation = 3.7mm). Seeds of the annual plants in this system germinate after rainstorms begin early in the winter, and plants complete their life cycle before the onset of the summer drought. The focal species of this experiment commonly grow together near outcrops of serpentine soil that are characterized by a low Ca:Mg ratio (Gram et al. 2004).

### Experiment Phase 1: Cultivating species-specific microbial communities

To cultivate the microbial community characteristic of each species’ soil, we grew five replicate high-density monocultures (8 g viable seed/m^2^) of each species in sterilized 3.6L pots. These pots were filled with 3L greenhouse soil (18.75% sand, 18.75% loam, 37.5% peat moss, 12.5% perlite, and 12.5% vermiculite) that we had autoclaved twice for 2 hours, with a 1-day rest period. To this sterile background we added 0.35L of field-collected inoculum, and capped this layer with 0.15L of sterilized greenhouse soil. This resulted in 10% v/v of live inoculum:sterile soil, a proportion that is consistent with other studies of plant-soil feedback (Crawford et al. 2019) and that minimizes abiotic differences among treatments. We collected the inoculum soil in Sedgwick reserve 1 week prior to the experiment and stored it at 0°C until planting. To ensure that the microbial community of this field soil was not pre-conditioned by any of the species in our experiment, we ensured that there were no individuals of our six focal species growing in a 1m radius around the five distinct points at which we collected the soil (the dominant plant around these points was the invasive grass *Avena fatua*).

We grew plants from seeds collected in Sedgwick reserve in the spring prior to the experiment. We surface-sterilized these seeds by soaking in 0.785% bleach for 3 minutes and washing in DI water twice for 1 minute each. After planting seeds, we stored the pots at 0°C for one week to trigger germination. Then, we allowed plants to grow in standard greenhouse conditions for 11 weeks, which is approximately the length of a complete growing season for these species. At the end of Phase 1, we harvested the aboveground biomass from each pot and homogenized the soil from the five replicate monocultures of each species to serve as the inoculum for the following phase of the experiment. We also saved soil samples from each replicate monoculture of *Plantago erecta* to use as the inoculum in a parallel experiment aimed to assess whether homogenizing across replicate cultivations influences the effects of cultivated soils on plant growth, described in Appendix S2.

### Experiment Phase 2: Quantifying plant responses to soil microbial communities

In the second phase of the experiment, we grew individuals of each of the six focal species in 125mL Deepots (Stuewe & Sons, Inc.) filled with 108mL greenhouse soil, autoclaved as for Phase 1, and 12mL of soil inoculum (again resulting in 10% v/v of live inoculum). The inoculum for each pot came from one of eight sources: a control treatment of autoclaved greenhouse soil, the same live field soil that was used to inoculate Phase 1 pots (and stored at 0°C during the first phase of the experiment), or soil cultivated by one of the six focal species during Phase 1 of the experiment (see Appendix S1 for graphical schematic of the experimental design). We grew 10 replicate individuals of each species in each soil background, for a total of 480 pots (6 species*8 soil sources*10 replicates), arranged in a randomized block design. We added three germinants of surface-sterilized seeds into each pot, and after 1 week thinned each pot to a single plant. We added an additional seedling in pots that had no surviving germinants after 1 week. Six pots had no surviving plants 2 weeks after initial planting; we excluded these pots from analyses. This phase of the experiment began in May 2019, and we harvested aboveground biomass after 8 weeks of growth, before the plants began senesce and lose aboveground biomass. We weighed the aboveground biomass of each individual after drying for 72H at 60°C.

### Data analysis: Quantifying pairwise stabilization and fitness differences

We used the log-transformed aboveground biomass of plants at the end of Phase 2 to calculate the degree of microbially mediated stabilization and fitness differences. Following Kandlikar et al. (2019), we calculated the degree to which plant-soil feedbacks stabilize each species pair by comparing growth growth in conspecific-cultivated soil community to growth in the heterospecific-cultivated community:

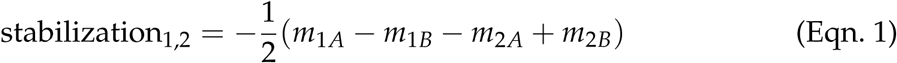

Each *m*_*ix*_ term represents the growth of plant species *i* with microbial community *x*, minus the plant species’ growth in the reference (uncultivated) field soil (e.g. *m*_1*A*_ = ln(biomass_sp 1, soil A_) − ln(biomass_sp 1, uncultivated soil_)). Due to arithmetic, plant growth on field soil cancels out of this equation and is not required to calculate microbially mediated stabilization (Bever et al. 1997, see Appendix S1). Positive values of stabilization indicate that soil microbes generate negative frequency dependent feedback loops that stabilize species interactions, whereas negative values indicate that plant-soil feedbacks drive positive frequency-dependent feedback loops that destabilize species interactions. The stabilization in Eqn. 1 is equal to negative one half of *I*_*S*_, the stabilization metric originally derived by Bever et al. (1997), and it allows for direct comparison with the microbially mediated fitness difference (Kandlikar et al. (2019); see Eqn. 3). This stabilization term is also equal to negative one half of the sum of the two species’ log response ratios (i.e.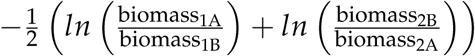, a metric commonly calculated in plant-soil feedback studies (Pernilla Brinkman et al. 2010, Crawford et al. 2019).

Inferring the net effects of plant-soil feedbacks requires also quantifying the microbially mediated fitness difference, which is calculated as the difference between the two species’ average response to cultivated soil microbial communities (Kandlikar et al. 2019):

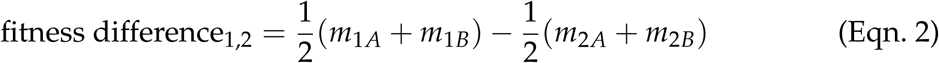

Although growth in the reference field soil is not required for calculating the degree of microbially mediated stabilization, this information *is* required for calculating the fitness difference. For simplicity, we always define species 1 in any given pair to be the fitness superior in this study.

Comparing the degree of microbially mediated stabilization and fitness difference allows us to infer the net effect of plant-soil feedbacks on species coexistence. Specifically, differences in plant responses to soil microbial communities result in species coexistence when stabilization is stronger than the microbially mediated fitness difference, or species exclusion when fitness differences mediated by microbes are larger than their stabilizing effects (Kandlikar et al. 2019):

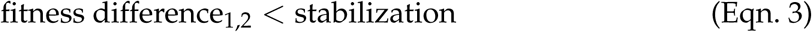

This condition in Eqn. 3 is equivalent to the feasibility criteria presented of Bever et al. (1997) and Eppinga et al. (2018), which states that plant-soil feedbacks allow coexistence provided that they cause a negative *I*_*S*_ *and* that the relative frequency of each species at equilibrium, calculated as 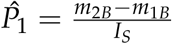 and 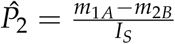, is between 0 and 1 (Appendix S3). To assess whether the microbially mediated fitness differences are weaker or stronger than the degree of stabilization, we calculated the stabilization and fitness difference metric within each replicate Phase 2 block, and summarized across blocks to calculate the mean and standard error.

We conducted all analyses in `R` v. 3.6.2 (R Core Team 2019) and provide code to recreate all analyses in Appendix S4. We will archive all data in an appropriate repository upon submission to a journal.

## Results

### Do the effects of cultivated soil microbial communities differ across species?

The influence of the soil microbial community on plant growth varied across plant species (Two-factor ANOVA, focal species x soil source interaction term *F*_35_ = 14.07, P < 0.001). Each plant species achieved its maximum biomass when growing with the field-collected reference soil, and five out of six focal species grew larger in sterile soils than when inoculated with soil cultivated by any species in Phase 1 (Fig. 1). Only one species (*Acmispon wrangelianus*) grew more poorly in sterile soil than in soil containing any live inoculum (88.9% lower biomass in sterile soil than average biomass with any live inoculum, Fig. 1).

**Figure 1:**
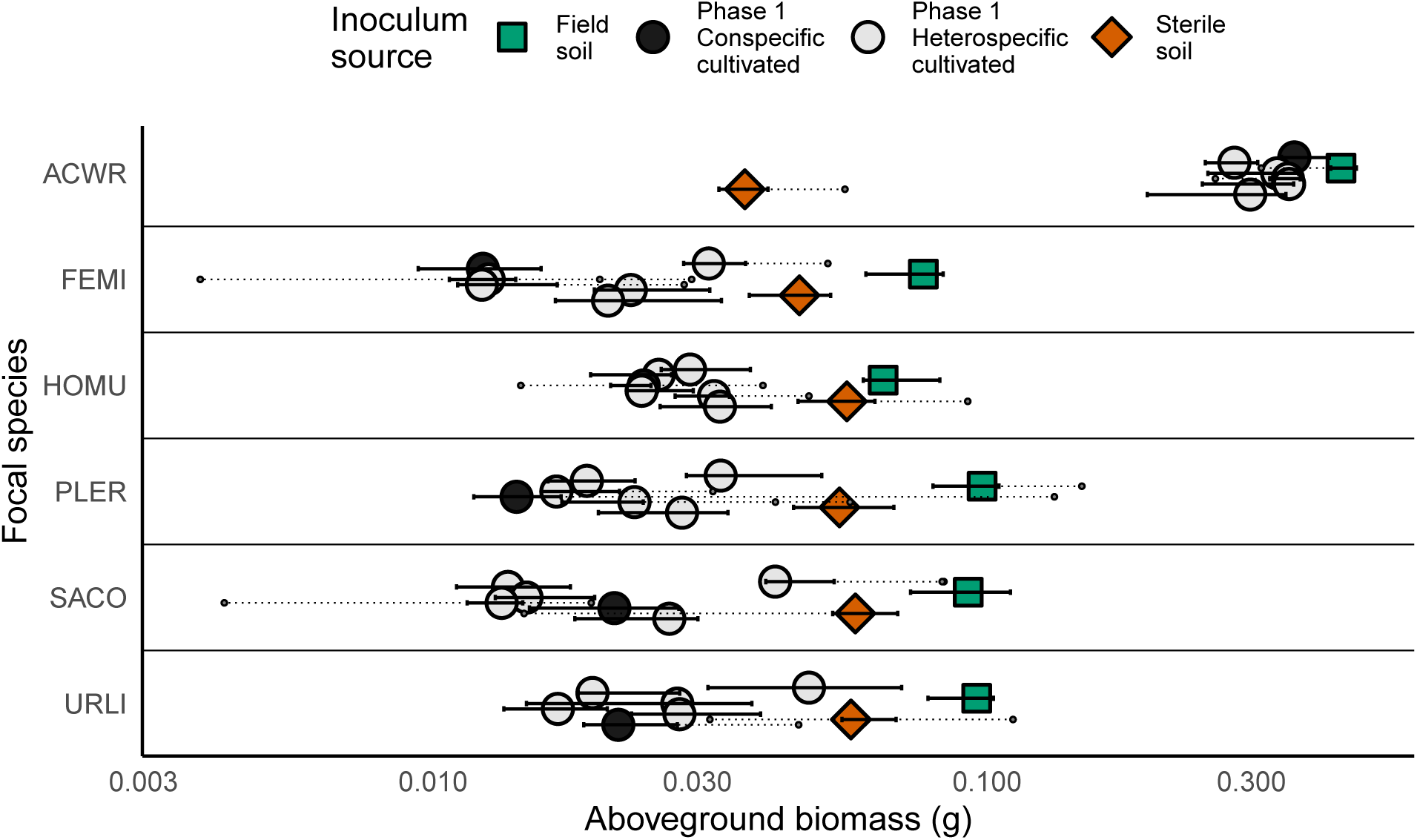
Effects of soil microbial inocula on plant growth. Aboveground biomass of each focal species growing with inocula of sterile greenhouse soil, live field-collected soil, or soil conditioned during phase 1. Large points indicate median biomass, and the solid error bars extend to the lower and upper quartiles. Small points and dashed lines show outliers, which were identified as points that were more than (1.5*IQR) away from the lower or upper quartile. Note the log-transformed X-axis.

### Do plant-soil feedbacks favor coexistence or exclusion among the focal species?

Plant species’ distinct responses to soil microbial communities result in complex coexistence outcomes for the species in this study. Plant-soil feedbacks tend to drive negative frequency-dependent feedback loops that stabilize the interaction for most (14/15) species pairs in our study (mean value of stabilization > 0), though species pairs differ in the strength of this effect (mean stabilization ± 2*SEM overlaps zero for 9 of these pairs). For a single species pair (*Salvia*/*Plantago*), microbial effects tend to drive positive frequency-dependent feedback loops that destabilize the pairwise interaction (mean value of stabilization < 0 but mean ± 2*SEM overlaps zero). Each species’ interaction with at least one other species is strongly stabilized by plant-soil feedbacks (Table 1).

**Table 1:** Microbially mediated stabilization and fitness differences among the fifteen species pairs in the study. Bold terms in the Stabilization and Fitness Difference columns indicate those values whose confidence intervals do not overlap zero. The net effect of plant-soil feedbacks reflects the relative magnitude of stabilization vs. fitness differences. When the stabilization is stronger than the microbially mediated fitness difference, the net effect of plant-soil feedbacks is to drive exclusion (first three rows). Alternately, plant-soil feedbacks drive exclusion when the fitness difference they mediate is larger than their stabilizing effect (bottom 12 rows). In six species pairs, there is especially strong evidence that plant-soil feedbacks drive exclusion, as the lower bound of the fitness difference estimate is larger than the upper bound of the stabilization estimate (final six rows).

| Species pair | Code | Stabilization | Fitness Difference | Net effect of PSF |
| --- | --- | --- | --- | --- |
| **Plant-soil feedbacks tend to promote coexistence (stabilization > fitness difference)** |  |  |  |  |
| *P. erecta*/ | PL_FE | 0.116 | 0.032 | coexistence |
| *F. microstachys* |  | (-0.13–0.362) | (-0.432–0.58) |  |
| *U. lindleyi*/ | UR_FE | 0.275 | 0.059 | coexistence |
| *F. microstachys* |  | (-0.029–0.579) | (-0.555–0.889) |  |
| *S. columbariae*/ | SA_UR | 0.161 | 0.112 | coexistence |
| *U. lindleyi* |  | (-0.147–0.469) | (-0.482–0.755) |  |
| **Plant-soil feedbacks tend to promote exclusion (fitness difference > stabilization)** |  |  |  |  |
| *A. wrangelianus*/ | AC_SA | **0.333** | **0.789** | exclusion |
| *S. columbariae* |  | (0.169–0.497) | (0.363–0.759) |  |
| *A. wrangelianus*/ | AC_UR | 0.227 | **0.834** | exclusion |
| *U. lindleyi* |  | (-0.069–0.523) | (0.468–0.593) |  |
| *F. microstachys*/ | FE_SA | 0.163 | 0.306 | exclusion |
| *S. columbariae* |  | (-0.027–0.353) | (-0.024–0.493) |  |
| *H. murinum*/ | HO_UR | **0.273** | **0.614** | exclusion |
| *U. lindleyi* |  | (0.055–0.491) | (0.182–0.705) |  |
| *P. erecta*/ | PL_UR | 0.123 | 0.173 | exclusion |
| *U. lindleyi* |  | (-0.207–0.453) | (-0.065–0.361) |  |
| *S. columbariae*/ | SA_PL | -0.105 | 0.024 | exclusion |
| *P. erecta* |  | (-0.453–0.243) | (-0.28–0.199) |  |
| **Strong evidence that plant-soil feedbacks promote exclusion (lower bound fitness difference estimate > upper bound of stabilization estimate)** |  |  |  |  |
| *A. wrangelianus*/ | AC_FE | **0.376** | **1.005** | exclusion |
| *F. microstachys* |  | (0.204–0.548) | (0.781–0.6) |  |
| *A. wrangelianus*/ | AC_HO | 0.045 | **0.724** | exclusion |
| *H. murinum* |  | (-0.053–0.143) | (0.57–0.199) |  |
| *A. wrangelianus*/ | AC_PL | **0.341** | **1.142** | exclusion |
| *P. erecta* |  | (0.077–0.605) | (0.846–0.637) |  |
| *H. murinum*/ | HO_FE | 0.031 | **0.689** | exclusion |
| *F. microstachys* |  | (-0.115–0.177) | (0.395–0.325) |  |
| *H. murinum*/ | HO_PL | **0.227** | **0.779** | exclusion |
| *P. erecta* |  | (0.017–0.437) | (0.451–0.555) |  |
| *H. murinum*/ | HO_SA | 0.038 | **0.613** | exclusion |
| *S. columbariae* |  | (-0.118–0.194) | (0.383–0.268) |  |

Soil microbes also drive a strong fitness difference in 9 out of the 15 species pairs in our study (mean fitness difference ± 2*SEM does not overlap zero). These microbially mediated fitness differences tend to favor certain species and harm others. In particular, the legume *Acmispon wrangelianus* gains a fitness advantage due to microbial feedbacks in its interactions with each of the other five species in our experiment. Similarly, the grass *Hordeum murinum* gains a fitness advantage over all species except *A. wrangelianus*.

Most importantly, using Eqn. 3 to compare the magnitude of microbially mediated stabilization and fitness differences reveals whether plant-soil feedbacks generally promote plant species coexistence or exclusion. We found that plant-soil feedbacks tend to promote species coexistence in three pairs in our study (*Plantago*/*Festuca, Salvia*/*Uropappus*, and *Uropappus/Festuca*). For these pairs, the mean stabilization estimate is larger than the mean microbially mediated fitness difference (Figure 2), though the confidence interval around the stabilization estimate overlaps that of the fitness difference (Table 1). However, for a majority of the species pairs in our study (12/15), larger microbially mediated fitness differences than stabilization suggest that plant-soil feedbacks tend to favor species exclusion over coexistence (Figure 2). There is especially strong evidence that soil microbes favor exclusion in six of these pairs, for which the lower bound of the fitness difference estimate is greater than the upper bound of the stabilization estimate (Table 1).

**Figure 2:**
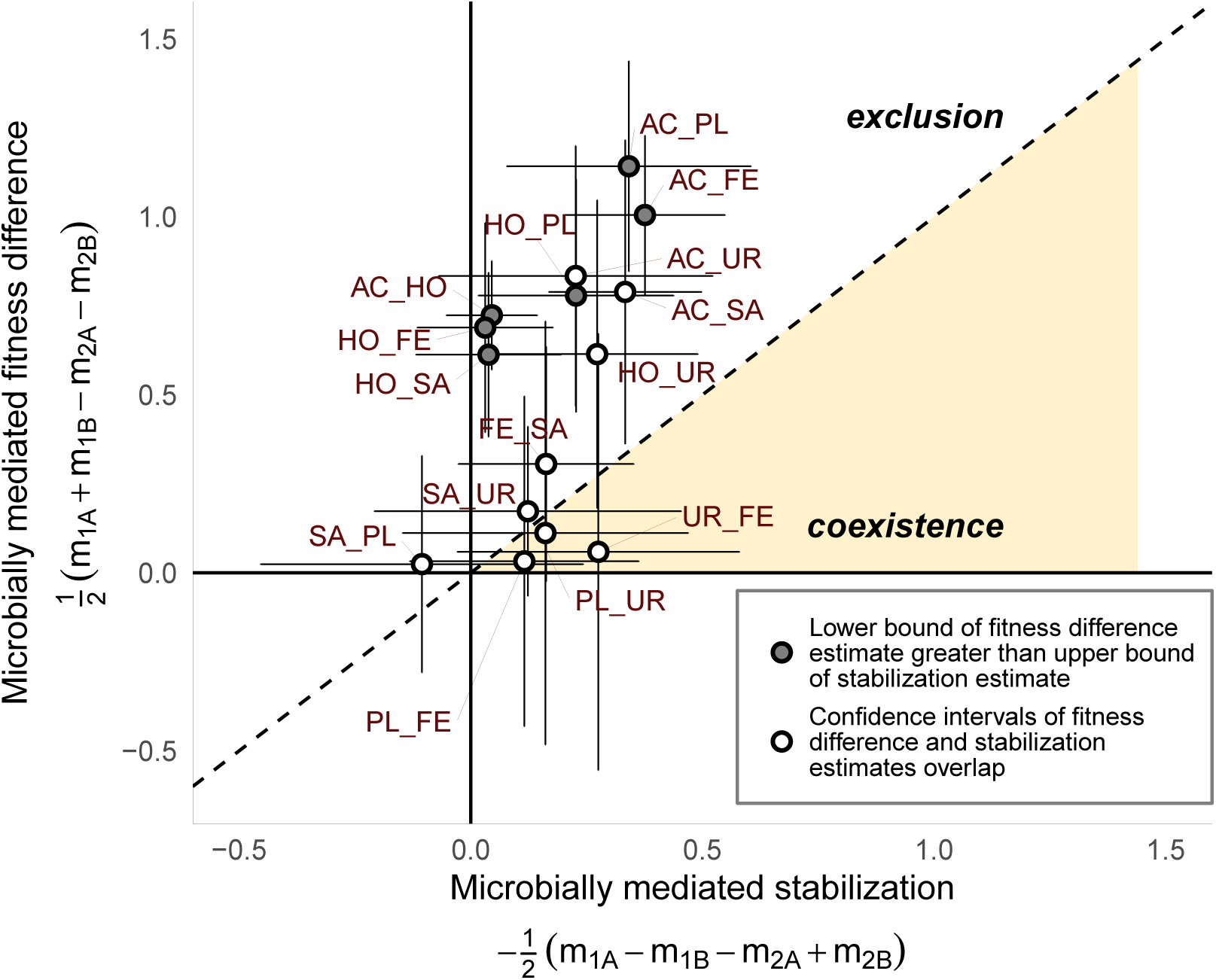
Stabilization and fitness differences calculated from the main experiment. Species pairs below the dashed line are predicted to coexist because the stabilizing effects of plant-soil feedbacks exceed the microbially mediated fitness difference, whereas microbially mediated fitness differences drive one species to exclusion in the pairs that fall above the dashed line. Error bars show mean ± 2*SEM.

## Discussion

Theoretical and empirical studies have shown that plant-soil feedbacks can influence whether a pair of species coexist if they drive stabilizing feedback loops that favor species that fall to low abundance and disadvantage more abundant species (or destabilizing loops that favor more abundant species and disadvantage rare ones) (Bever et al. 1997, Crawford et al. 2019). However, recent theoretical advances have clarified that species coexistence is also determined by the degree to which soil micobes mediate an average fitness difference that gives one species a demographic advantage over its competitor, regardless of its abundance in the system (Chesson 2000, Kandlikar et al. 2019, Ke and Wan 2019). Here, we show that empirically quantifying the microbially mediated fitness difference and comparing it to the degree of microbial stabilization in Bever et al. (1997)’s classic model of plant-soil feedback is essential for understanding how soil microbes shape plant species coexistence. Specifically, we found that even though soil microbes stabilize pairwise interactions by generating negative frequency-dependent dynamics in nearly all pairs in our study, their overall effect is to often favor species exclusion because they mediate substantial fitness differences.

That plant-soil feedbacks drive stabilization but also strong average fitness differences is determined by the particular arrangement of interspecific differences in species’ response to soil communities cultivated by conspecifics and heterospecifics. In our experiment, species generally performed better with a soil community cultivated by heterospecifics than with a soil community cultivated by conspecifics (across all species, growth with conspecific-cultivated microbes was on average 13% lower than growth with heterospecific microbes). Thus, the effect of plant-soil feedbacks is to generally stabilize rather than destabilize plant interactions (Fig. 2), a result which is consistent with meta-analyses of similar experiments among grassland species (Kulmatiski et al. 2008, Crawford et al. 2019). Inferring the influence of plant-soil feedbacks from this result alone might lead us to conclude that soil microbes generally favor plant diversity in this system. However, a novel aspect of our experiment is that we can compare the stabilizing effects of soil microbes to the fitness difference they generate to more fully assess their effects on plant coexistence. This comparison is possible because we also measured plant growth with the microbial community of field-collected soils whose microbial community has not been directly influenced of any focal species in our experiment, a treatment has been omitted from most studies of plant-soil feedbacks.

All six species in our experiment grew less vigorously with microbes cultivated during Phase 1 than when grown with the microbial community of uncultivated field soil (Fig. 1). This indicates that greenhouse-grown high-density monocultures of these species may harbor more pathogenic (or fewer mutualistic) soil microbes than field-collected reference soils. However, this effect was weak for *Acmispon wrangelianus* – the only Fabaceae species in our experiment – which grew more similarly with microbes cultivated during Phase 1 as with the field-collected reference microbial community (*A. wrangelianus* growth was 28% lower with cultivated microbes than with the reference field inoculum, vs. 71% lower growth on average for all other species). This result, as well as our finding that *A. wrangelianus* grows much more vigorously when inoculated with any live microbial community than in sterile soil (Fig. 1), is consistent with other studies showing that growth of *A. wrangelianus* benefits from the presence of many strains of nitrogen-fixing soil bacteria in the genus *Mesorhizobium*, strains likely present in our field soil inoculum (Porter et al. 2016, 2019). As a consequence of the differences in species’ response to the microbial communities cultivated by the focal species, plant-soil feedbacks generate a fitness advantage in favor of *A. wrangelianus* in its pairwise interaction with each of the other species in our experiment (Table 1). Although empirical studies of plant-soil feedback have rarely explicitly quantified microbially mediated average fitness differences, our is one of a growing number of studies finding that complex, diverse soil microbial communities benefit Fabaceae species that associate with N-fixing bacteria more than plants of other functional groups (van der Heijden et al. 2015, Teste et al. 2017). This indicates that plant-soil feedbacks may frequently drive a fitness advantage in favor of these legume species.

We quantified the growth of plants with soil microbes cultivated by conspecifics and heterospecifics, as well as growth with an reference microbial community from field-collected soil. This design isolates the effect of soil microbes on plant interactions, but more thoroughly evaluating the consequences of soil microbes on plant diversity requires also considering other processes like resource competition that simultaneously influence species interactions (Bever 2003, Kandlikar et al. 2019). Specifically, although stabilizing plant-soil feedbacks promote species coexistence regardless of other processes, the microbially mediated fitness differences that drive exclusion in Bever et al. (1997)’s model may in fact favor plant diversity in nature if they benefit the otherwise weaker resource competitor. Indeed, in a previous field experiment among annual plants in the same system that motivated our study, *A. wrangelianus* was predicted to be excluded in pairwise interactions with three other focal species in our experiment (*Plantago erecta, Salvia columbariae*, and *Uropappus lindelyi*) (Kraft et al. 2015, note that *U. lindelyi* was called *Agoseris heterophylla* in that study). In that study, *A. wrangelianus* was grown with competitors in live field soil, so the results reflect the joint effects of resource competition, plant-soil feedback, and other processes operating in nature. Thus, the microbially mediated fitness advantage in favor of *A. wrangelianus* that we identified appears to simply improve the performance of a weak competitor, though the relative boost to *A. wrangelianus’* performance due to the microbially mediated fitness advantage is not sufficient to overcome other competitive asymmetries. More generally, if soil microbes drive observed trade-offs between plant species’ competitive ability and sensitivity to pathogenic or mutualistic microbes (Bever et al. 2015, Peay 2016), microbially mediated fitness differences might often promote plant diversity by reducing the degree of niche differentiation required for stable coexistence. Future empirical studies designed to simultaneously quantify the effects of competitors and soil microbes (e.g. using the designs proposed by Ke and Wan (2019)) will help clarify the interplay between plant-soil feedback and resource competition.

Our study highlights the potential for plant-soil feedbacks to simultaneously stabilize pairwise interactions and also drive average fitness differences that always favor one species over the other, but our results have some important caveats. First, we mixed the soils cultivated by replicate Phase 1 monocultures of each species to create the soil inoculum for Phase 2. This is a common step in many plant-soil feedback studies (e.g. Klinerová and Dostál (2020); Cortois et al. (2016)), but it can result in falsely precise or inflated estimates of the soil microbial community’s effect on plant growth (Reinhart and Rinella 2016). When we assessed the consequence of soil homogenization by comparing growth in soil homogenized across replicate monocultures vs. growth with soil from a single monoculture, we found that this homogenizing across monocultures is unlikely to have significantly influenced the results of our main experiment (Fig. S2.1). In general, however, we agree with Reinhart and Rinella (2016) that more careful study of the variable nature of plant-microbe interactions will be a fruitful avenue for future research. A second limitation of our study is that it did not account for the fact that the composition and dynamics of soil microorganisms in Mediterranean ecosystems are also influenced by the length and intensity of the summer drought that separates the annual plant community’s growing seasons (Barnard et al. 2014). By not accouting for the possibility that the cultivating effects of plant species on soil microbial communities erode over the six-month dry season, our experiment might have overestimated the effects of plant-soil feedbacks on species coexistence in this annual plant system. Future studies that adapt the standard two-phase design of plant-soil feedback experiments to capture the biological idiosyncrasies of the focal communities will be important for contextualizing our understanding of how soil microorganisms shape diversity in natural plant communities (Smith-Ramesh and Reynolds 2017).

In conclusion, we have demonstrated that dynamic feedbacks between plants and soil microorganisms can have important consequences on plant coexistence. Existing research has emphasized the potential for such feedbacks to stabilize or destabilize plant interactions by generating negative or positive frequency-dependent dynamics. Here we highlight that inferring the net effects of soil microbes by evaluating only their (de)stabilizing effects and not comparing these to the microbially mediated fitness difference can lead to false conclusions regarding soil microbes’ effects on plant diversity. Moving forward, translating the consequences of microbially mediated stabilization and fitness difference into predictions for how soil microbes mediate diversity in nature will require contextualizing the effects of soil microbes relative to those of competition and other processes that affect the dynamics of plant and soil microbial communities.

## Acknowledgements

We acknowledge the Tongva/Gabriellino and Chumash peoples as the traditional land caretakers of the ecosystem studied in this project. We thank Anmol Dhaliwal, Jonathan Shi, and the UCLA Plant Growth Facility staff for help with the greenhouse experiment, and Kate McCurdy and other staff at Sedgwick Reserve for help in the field. For comments on early versions of the manuscript, we thank Madeline Cowen, Kenji Hayashi, Andy Kleinhesselink, Mary Van Dyke, and Marcel Vaz. This work was funded by the American Naturalist Society Student Research Award and the La Kretz Center for Conservation Science. GSK was supported by the National Science Foundation Graduate Research Fellowship (DGE-1650604) and by the UCLA Dept. of Ecology and Evolutionary Biology, XY was supported by the CALeDNA Summer Undergraduate Internship, and NJBK and JML were supported by the National Science Foundation DEB-1644641.

