## Appendix for "Quantifying microbially mediated fitness differences reveals the tendency for plant-soil feedbacks to drive species exclusion among California annual plants"

Last rendered 13 Feb 2020

### Appendix S1: Modifying the standard two-phase design of plant-soil feedback experiments to quantify microbially mediated fitness differences

Most studies that use a two-phase experimental design to study the coexistence consequences of plant-soil feedbacks only grow plants with cultivated soil microbial communities in the second phase of their experiment, and not with an uncultivated, reference soil community. This is because quantifying microbially mediated stabilization, which is the focus of most such studies, does not require measuring plant growth with uncultivated microbes. Specifically, Bever et al. (1997) showed that following the original definitions of the $m_{ix}$ terms as growth of plant $i$ with microbes $x$ minus the growth of plant $i$ with uncultivated microbes ($m_{ix} = G_{ix} - G_{iO}$), growth with uncultivated microbes is irrelavent for quantifying the stabilization metric $I_S$. Following the same logic, growth with uncultivated microbial communities also cancels out of the equation for the stabilization metric derived in Kandlikar et al. (2019):

$$\begin{aligned} \text{stabilization} &= -\frac{1}{2}(m_{1A} + m_{2B} - m_{1B} - m_{2A}) \\ &= -\frac{1}{2}((G_{1A} - \cancel{G_{1O}}) + (G_{2B} - \cancel{G_{2O}}) - (G_{1B} - \cancel{G_{1O}}) - (G_{2A} - \cancel{G_{2O}})) \\ &= -\frac{1}{2}(G_{1A} + G_{2B} - G_{1B} - G_{2A}) \end{aligned} \tag{1}$$

However, growth with uncultivated microbes *is* relevant for quantifying the microbially mediated fitness difference:

$$\begin{aligned} \text{fitness difference}_{1,2} &= \frac{1}{2}(m_{1A} + m_{1B}) - \frac{1}{2}(m_{2A} + m_{2B}) \\ &= \frac{1}{2}((G_{1A} - G_{1O}) + (G_{1B} - G_{1O})) - \frac{1}{2}((G_{2A} - G_{2O}) + (G_{2B} - G_{2O})) \\ &= \left(\frac{1}{2}(G_{1A} + G_{1B}) - G_{1O}\right) - \left(\frac{1}{2}(G_{2A} + G_{2B}) - G_{2O}\right) \end{aligned} \tag{2}$$

Thus, simply using the standard two-phase design of plant soil feedback experiments with an additional treatment of growth in uncultivated reference soil in the second phase (see Fig. S1.1), gives all the measures required to quantify both the stabilization and fitness difference mediated by soil microbes in Bever et al. (1997)'s model of plant-soil feedback.

**Figure S1.1: Schematic of experimental design.** Note that plants were grown as high-density monocultures (8g seed/m2) for the first (cultivation) phase, but shown as a single plant here for simplicity. Plants were grown as single individuals in pots for the second (response) phase of the experiment. Soils across the five replicate Phase 1 cultivations were homogenized to create the inocula for the second phase of the experiment (but see *Variation experiment* and Fig. 3 of the main text for results from a parallel experiment in which we tested the effects of each *Plantago*-cultivated soil separately). Our experiment included six focal species; we show only two here for clarity.

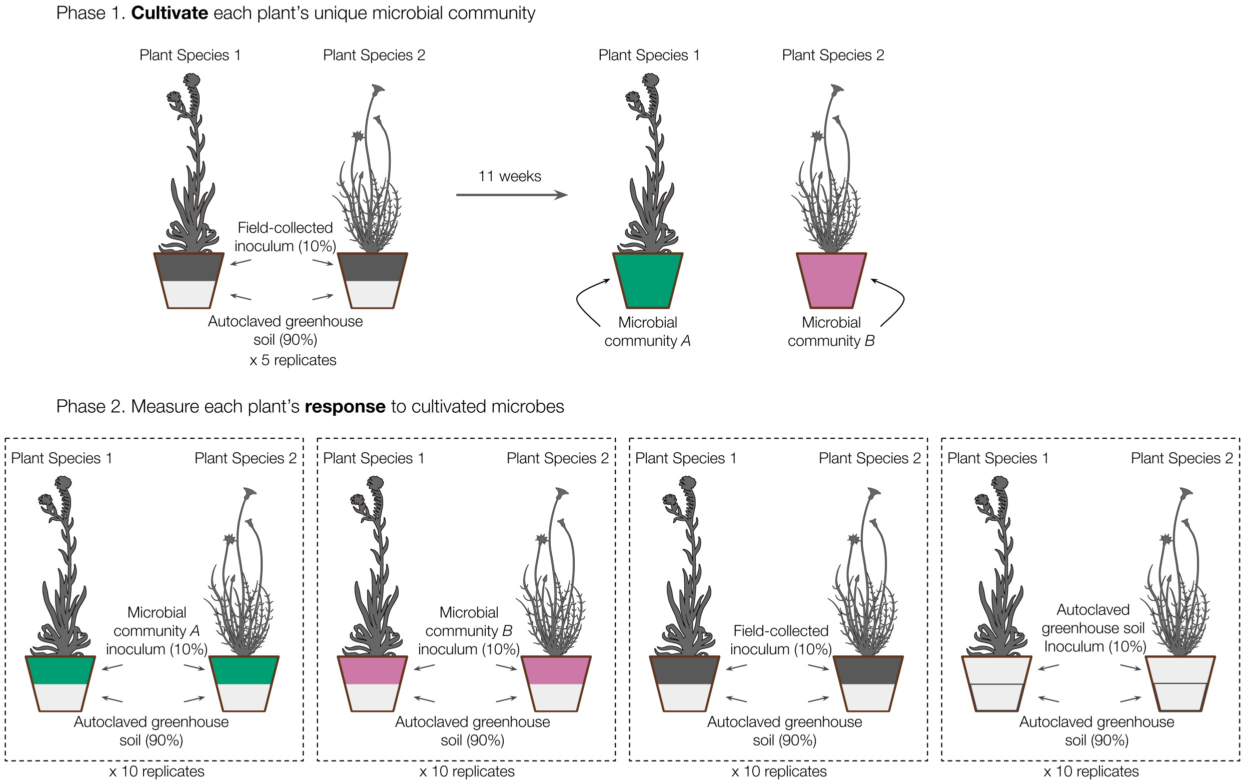

### Appendix S2: Evaluating the effects of homogenizing replicate Phase 1 cultivations

Homogenizing soils from across the replicate Phase 1 cultivations to create the inocula for the second phase, a common step in plant-soil feedback experiments (Gundale et al. 2018) that we adopt in this study, makes such experiments more feasible but can lead to biased and falsely precise results (Reinhart and Rinella 2016). To explore the variation in the effects of the replicate Phase 1 cultivations, we set up a parallel experiment in which we grew ten replicate individuals of *Plantago erecta* and *Festuca microstachys* in soils cultivated by each of the five *Plantago* Phase 1 monocultures (2 species * 5 soil inocula * 10 replicates = 100 pots). Although this approach does not quantify the variation in each focal species' cultivation, it can yield insight into the potential consequences of pooling across replicate Phase 1 monocultures on plant growth. As in Phase 2 of the feedback experiment, we planted germinants from surface-sterilized seeds into 125mL Deepots that contained 10% v/v live inoculum, and added an additional seedling in pots that had no surviving germinants after 1 week. Two pots had no surviving plants 2 weeks after initial planting; we excluded these pots from analyses. We harvested aboveground biomass after 8 weeks of growth, and weighed after drying for 72H at 60°C.

**Results:** The biomass of *Plantago erecta* and *Festuca microstachys* grown with an inoculum that came from a single *Plantago* phase 1 monoculture did not differ significantly from their biomass when grown with an inoculum made by combining soil from each of the five *Plantago* monocultures (one-factor Anova; Plantago: $F_{5,55} = 1.16$, $p > 0.05$; Festuca: $F_{5,52} = 1.78$, $p > 0.05$, Fig. S2.1). Whether the inoculum was sourced from a single Phase 1 monoculture or was homogenized across replicate monocultures also did not influence the variance in aboveground biomass (Levene's test; Plantago $F_{5,55} = 0.82$, $p > 0.05$; Festuca $F_{5,52} = 0.44$; $p > 0.05$).

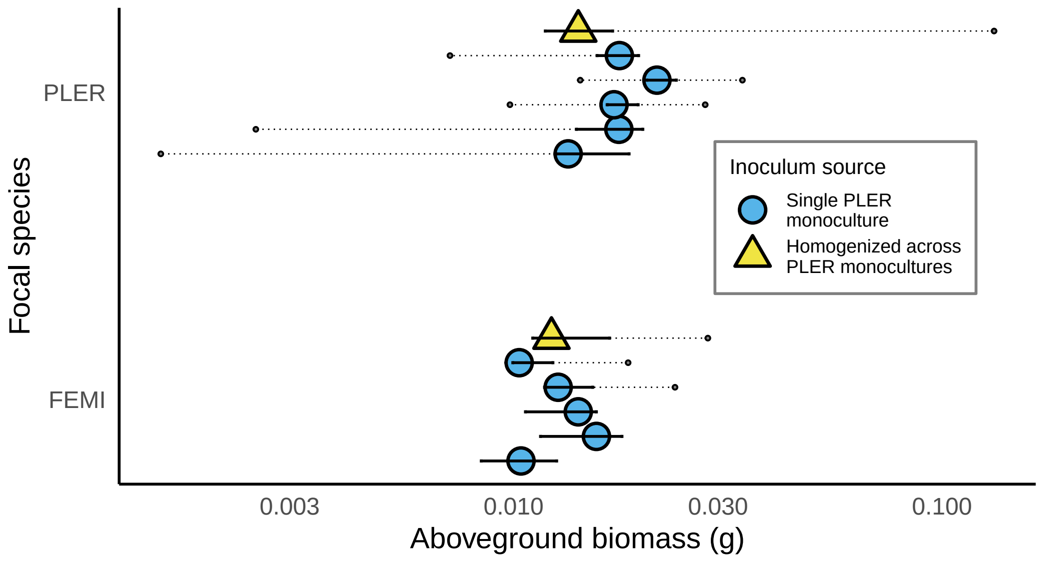

**Figure S2.1: Results from a parallel experiment to explore the effect of homogenizing soil from replicate Phase 1 monocultures on growth of two plant species.** Aboveground biomass of *Plantago erecta* and *Festuca microstachys* grown with a soil inoculum that was sourced either from a single *Plantago* phase 1 monoculture (blue circles) or with an inoculum created by homogenizing soil from all five repilicate monocultures (yellow triangles), as was done for the main experiment. Large points indicate median biomass, and the solid error bars extend to the lower and upper quartiles. Small points and dashed lines show outliers, which were identified as points that were more than (1.5*IQR) away from the lower or upper quartile. Note the log-transformed X-axis.

### Appendix S3: Comparing predictions of coexistence derived by comparing strength of stabilization and fitness differences to predictions made via Bever et al. (1997)'s feasibility analysis

Our approach of inferring coexistence in terms of microbially mediated stabilization and fitness differences using our experimental data yields conclusions that are consistent with the feasibility analysis originally presented in Bever et al. (1997) (for an algebraic explanation of this equivalence, see Appeddix S1 of Kandlikar et al. (2019)). In Bever et al. (1997)'s analysis of the plant-soil feedback model, soil microbes stabilize plant interactions that they result in a negative value for the stabilization metric $I_S$, calculated as $I_S = m_{1A} + m_{2B} - m_{1B} - m_{2A}$. Importantly, negative $I_S$ is a *necessary* condition for plant coexistence, but it is not *sufficient*: stable coexistence of plant species also requires that both plants have a positive frequency at equilibrium, with this equilibrium frequency $\hat{P}$ calculated as $\hat{P}_1 = \frac{m_{2B} - m_{1B}}{I_S}$ for species 1, and $\hat{P}_2 = \frac{m_{1A} - m_{2A}}{I_S}$ for species 2. In other words, plant-soil feedbacks result in stable coexistence provided three conditions are satisfied (Bever et al. 1997, Eppinga et al. 2018):

$$I_S < 0; \quad 0 < \hat{P}_1 < 1; \quad 0 < \hat{P}_2 < 1 \qquad \text{(Eqn. S1.2)}$$

In Table S1.1 below we show, for each species pair in each block, that analyzing experimental data in terms of Eqn S1.1 or Eqn S1.2 yields the same conclusions regarding coexistence vs. exclusion.

| rep | pair | $m_{1A}$ | $m_{1B}$ | $m_{2A}$ | $m_{2B}$ | $I_S$ | $\hat{p}_1$ | $\hat{p}_2$ | outcome (feasibility) | stabilization | fitness difference | outcome (IGR) |
|---|---|---|---|---|---|---|---|---|---|---|---|---|
| 1 | AC_FE | -0.410 | -0.219 | -0.721 | -1.720 | -1.191 | 1.260 | -0.260 | exclude | 0.596 | 0.906 | exclude |
| 2 | AC_FE | -0.242 | -0.346 | -1.035 | -2.290 | -1.151 | 1.689 | -0.689 | exclude | 0.575 | 1.369 | exclude |
| 3 | AC_FE | -0.193 | -0.743 | -1.155 | -2.224 | -0.519 | 2.854 | -1.854 | exclude | 0.259 | 1.222 | exclude |
| 4 | AC_FE | -0.282 | -0.275 | -0.687 | -1.628 | -0.949 | 1.426 | -0.426 | exclude | 0.475 | 0.879 | exclude |
| 5 | AC_FE | -0.231 | -0.488 | -1.060 | -1.914 | -0.597 | 2.387 | -1.387 | exclude | 0.299 | 1.127 | exclude |
| 6 | AC_FE | -0.186 | -0.360 | -0.604 | -2.499 | -1.721 | 1.243 | -0.243 | exclude | 0.861 | 1.278 | exclude |
| 7 | AC_FE | -0.165 | -0.667 | -0.842 | -1.901 | -0.557 | 2.215 | -1.215 | exclude | 0.278 | 0.955 | exclude |
| 8 | AC_FE | 0.447 | -0.390 | -1.117 | -1.637 | 0.316 | -3.942 | 4.942 | exclude | -0.158 | 1.405 | exclude |
| 9 | AC_FE | 0.052 | -0.645 | -0.012 | -1.146 | -0.437 | 1.146 | -0.146 | exclude | 0.218 | 0.282 | exclude |
| 10 | AC_FE | -0.280 | -0.521 | -0.552 | -1.506 | -0.713 | 1.382 | -0.382 | exclude | 0.356 | 0.629 | exclude |
| 1 | AC_HO | -0.410 | -0.532 | -1.133 | -0.887 | 0.368 | -0.965 | 1.965 | exclude | -0.184 | 0.539 | exclude |
| 2 | AC_HO | -0.242 | -0.171 | -0.976 | -1.088 | -0.183 | 5.003 | -4.003 | exclude | 0.092 | 0.826 | exclude |
| 3 | AC_HO | -0.193 | -0.849 | -0.665 | -1.463 | -0.142 | 4.307 | -3.307 | exclude | 0.071 | 0.542 | exclude |
| 4 | AC_HO | -0.282 | -0.016 | -0.724 | -0.843 | -0.385 | 2.147 | -1.147 | exclude | 0.193 | 0.635 | exclude |
| 5 | AC_HO | -0.231 | -0.398 | -1.144 | -1.570 | -0.260 | 4.515 | -3.515 | exclude | 0.130 | 1.042 | exclude |
| 6 | AC_HO | -0.186 | -0.276 | -0.749 | -1.138 | -0.299 | 2.879 | -1.879 | exclude | 0.150 | 0.712 | exclude |
| 7 | AC_HO | -0.165 | -0.820 | -0.711 | -0.955 | 0.411 | -0.329 | 1.329 | exclude | -0.205 | 0.341 | exclude |
| 8 | AC_HO | 0.447 | 0.131 | -0.829 | -0.914 | 0.232 | -4.506 | 5.506 | exclude | -0.116 | 1.160 | exclude |
| 9 | AC_HO | 0.052 | -0.325 | -0.592 | -1.192 | -0.223 | 3.892 | -2.892 | exclude | 0.111 | 0.755 | exclude |
| 10 | AC_HO | -0.280 | -0.204 | -0.750 | -1.101 | -0.427 | 2.101 | -1.101 | exclude | 0.213 | 0.683 | exclude |
| 1 | AC_PL | -0.410 | -0.375 | -0.795 | -1.422 | -0.662 | 1.582 | -0.582 | exclude | 0.331 | 0.716 | exclude |
| 2 | AC_PL | -0.242 | -0.165 | -1.225 | 0.324 | 1.472 | 0.332 | 0.668 | exclude | -0.736 | 0.247 | exclude |
| 3 | AC_PL | -0.193 | -0.213 | -0.887 | -1.486 | -0.579 | 2.198 | -1.198 | exclude | 0.290 | 0.983 | exclude |
| 4 | AC_PL | -0.282 | -0.343 | -1.321 | -2.242 | -0.862 | 2.205 | -1.205 | exclude | 0.431 | 1.469 | exclude |
| 5 | AC_PL | -0.231 | -0.094 | -0.959 | -1.570 | -0.748 | 1.972 | -0.972 | exclude | 0.374 | 1.102 | exclude |
| 6 | AC_PL | -0.186 | -0.164 | -1.166 | -1.843 | -0.700 | 2.399 | -1.399 | exclude | 0.350 | 1.330 | exclude |
| 7 | AC_PL | -0.165 | -0.361 | -0.685 | -2.069 | -1.188 | 1.437 | -0.437 | exclude | 0.594 | 1.114 | exclude |
| 8 | AC_PL | 0.447 | 0.076 | -1.262 | -2.291 | -0.659 | 3.594 | -2.594 | exclude | 0.329 | 2.038 | exclude |
| 9 | AC_PL | 0.052 | -0.251 | -0.322 | -2.326 | -1.701 | 1.220 | -0.220 | exclude | 0.850 | 1.224 | exclude |
| 10 | AC_PL | -0.280 | -0.688 | -0.881 | -2.485 | -1.197 | 1.502 | -0.502 | exclude | 0.598 | 1.199 | exclude |
| 1 | AC_SA | -0.410 | -0.147 | -0.237 | -0.915 | -0.942 | 0.816 | 0.184 | coex | 0.471 | 0.297 | coex |
| 3 | AC_SA | -0.193 | -0.642 | -0.842 | -1.961 | -0.670 | 1.969 | -0.969 | exclude | 0.335 | 0.984 | exclude |
| 4 | AC_SA | -0.282 | -0.895 | -0.455 | -1.112 | -0.045 | 4.820 | -3.820 | exclude | 0.023 | 0.195 | exclude |
| 5 | AC_SA | -0.231 | -0.166 | -1.117 | -1.421 | -0.369 | 3.397 | -2.397 | exclude | 0.185 | 1.070 | exclude |

| | | | | | | | | | | | | |
|---|---|---|---|---|---|---|---|---|---|---|---|---|
| 7 | AC_SA | -0.165 | -0.332 | 0.107 | -1.624 | -1.565 | 0.826 | 0.174 | coex | 0.782 | 0.510 | coex |
| 8 | AC_SA | 0.447 | 0.157 | -1.409 | -2.156 | -0.458 | 5.056 | -4.056 | exclude | 0.229 | 2.085 | exclude |
| 9 | AC_SA | 0.052 | -0.717 | -0.421 | -1.439 | -0.249 | 2.903 | -1.903 | exclude | 0.124 | 0.598 | exclude |
| 10 | AC_SA | -0.280 | -0.191 | -0.474 | -1.144 | -0.760 | 1.255 | -0.255 | exclude | 0.380 | 0.573 | exclude |
| 1 | AC_UR | -0.410 | -0.320 | -1.118 | -1.695 | -0.668 | 2.061 | -1.061 | exclude | 0.334 | 1.042 | exclude |
| 2 | AC_UR | -0.242 | -0.382 | -0.093 | -1.189 | -0.956 | 0.844 | 0.156 | coex | 0.478 | 0.329 | coex |
| 4 | AC_UR | -0.282 | -0.855 | -0.875 | -2.639 | -1.191 | 1.498 | -0.498 | exclude | 0.596 | 1.189 | exclude |
| 5 | AC_UR | -0.231 | -1.003 | -0.295 | -1.480 | -0.414 | 1.155 | -0.155 | exclude | 0.207 | 0.271 | exclude |
| 6 | AC_UR | -0.186 | -0.206 | -0.935 | -2.162 | -1.207 | 1.620 | -0.620 | exclude | 0.604 | 1.352 | exclude |
| 7 | AC_UR | -0.165 | -0.269 | -2.090 | -1.314 | 0.880 | -1.188 | 2.188 | exclude | -0.440 | 1.485 | exclude |
| 8 | AC_UR | 0.447 | -1.386 | -0.330 | -0.956 | 1.206 | 0.356 | 0.644 | exclude | -0.603 | 0.174 | exclude |
| 9 | AC_UR | 0.052 | -0.224 | -0.390 | -1.441 | -0.775 | 1.570 | -0.570 | exclude | 0.388 | 0.830 | exclude |
| 1 | FE_HO | -1.720 | -1.456 | -0.782 | -0.887 | -0.369 | -1.542 | 2.542 | exclude | 0.185 | -0.754 | exclude |
| 2 | FE_HO | -2.290 | -1.844 | -1.190 | -1.088 | -0.344 | -2.196 | 3.196 | exclude | 0.172 | -0.928 | exclude |
| 3 | FE_HO | -2.224 | -2.023 | -0.842 | -1.463 | -0.822 | -0.682 | 1.682 | exclude | 0.411 | -0.971 | exclude |
| 4 | FE_HO | -1.628 | -1.537 | -1.395 | -0.843 | 0.461 | 1.505 | -0.505 | exclude | -0.231 | -0.464 | exclude |
| 5 | FE_HO | -1.914 | -2.039 | -1.066 | -1.570 | -0.380 | -1.233 | 2.233 | exclude | 0.190 | -0.658 | exclude |
| 6 | FE_HO | -2.499 | -2.690 | -1.375 | -1.138 | 0.428 | 3.628 | -2.628 | exclude | -0.214 | -1.338 | exclude |
| 7 | FE_HO | -1.901 | -1.805 | -1.103 | -0.955 | 0.051 | 16.530 | -15.530 | exclude | -0.026 | -0.824 | exclude |
| 8 | FE_HO | -1.637 | -1.676 | -0.918 | -0.914 | 0.043 | 17.627 | -16.627 | exclude | -0.022 | -0.741 | exclude |
| 9 | FE_HO | -1.146 | -0.577 | -1.444 | -1.192 | -0.316 | 1.946 | -0.946 | exclude | 0.158 | 0.457 | exclude |
| 10 | FE_HO | -1.506 | -2.082 | -1.149 | -1.101 | 0.624 | 1.571 | -0.571 | exclude | -0.312 | -0.669 | exclude |
| 1 | FE_PL | -1.720 | -1.656 | -1.506 | -1.422 | 0.020 | 11.631 | -10.631 | exclude | -0.010 | -0.224 | exclude |
| 2 | FE_PL | -2.290 | -1.966 | -1.801 | 0.324 | 1.800 | 1.272 | -0.272 | exclude | -0.900 | -1.389 | exclude |
| 3 | FE_PL | -2.224 | -1.856 | -1.335 | -1.486 | -0.519 | -0.714 | 1.714 | exclude | 0.259 | -0.630 | exclude |
| 4 | FE_PL | -1.628 | -1.288 | -2.260 | -2.242 | -0.322 | 2.965 | -1.965 | exclude | 0.161 | 0.793 | exclude |
| 5 | FE_PL | -1.914 | -2.020 | -1.251 | -1.570 | -0.214 | -2.106 | 3.106 | exclude | 0.107 | -0.557 | exclude |
| 6 | FE_PL | -2.499 | -1.583 | -2.160 | -1.843 | -0.600 | 0.434 | 0.566 | coex | 0.300 | -0.040 | coex |
| 7 | FE_PL | -1.901 | -1.826 | -1.508 | -2.069 | -0.635 | 0.383 | 0.617 | coex | 0.318 | -0.075 | coex |
| 8 | FE_PL | -1.637 | -1.749 | -1.660 | -2.291 | -0.519 | 1.044 | -0.044 | exclude | 0.260 | 0.283 | exclude |
| 9 | FE_PL | -1.146 | -0.609 | -1.723 | -2.326 | -1.139 | 1.507 | -0.507 | exclude | 0.570 | 1.147 | exclude |
| 10 | FE_PL | -1.506 | -2.026 | -1.779 | -2.485 | -0.186 | 2.466 | -1.466 | exclude | 0.093 | 0.366 | exclude |
| 1 | FE_SA | -1.720 | -0.827 | -1.043 | -0.915 | -0.766 | 0.116 | 0.884 | coex | 0.383 | -0.294 | coex |
| 3 | FE_SA | -2.224 | -1.377 | -1.848 | -1.961 | -0.960 | 0.608 | 0.392 | coex | 0.480 | 0.104 | coex |
| 4 | FE_SA | -1.628 | -1.151 | -2.361 | -1.112 | 0.771 | 0.051 | 0.949 | exclude | -0.386 | 0.347 | exclude |

| | | | | | | | | | | | | |
|---|---|---|---|---|---|---|---|---|---|---|---|---|
| 5 | FE_SA | -1.914 | -1.201 | -2.142 | -1.421 | 0.007 | -29.649 | 30.649 | exclude | -0.004 | 0.224 | exclude |
| 7 | FE_SA | -1.901 | -1.698 | -1.930 | -1.624 | 0.103 | 0.714 | 0.286 | exclude | -0.052 | -0.022 | exclude |
| 8 | FE_SA | -1.637 | -1.042 | -2.137 | -2.156 | -0.614 | 1.814 | -0.814 | exclude | 0.307 | 0.807 | exclude |
| 9 | FE_SA | -1.146 | -0.391 | -2.407 | -1.439 | 0.213 | -4.917 | 5.917 | exclude | -0.107 | 1.154 | exclude |
| 10 | FE_SA | -1.506 | -0.912 | -1.527 | -1.144 | -0.211 | 1.101 | -0.101 | exclude | 0.105 | 0.127 | exclude |
| 1 | FE_UR | -1.720 | -0.543 | -1.864 | -1.695 | -1.009 | 1.143 | -0.143 | exclude | 0.504 | 0.648 | exclude |
| 2 | FE_UR | -2.290 | -3.403 | -0.983 | -1.189 | 0.907 | 2.440 | -1.440 | exclude | -0.454 | -1.760 | exclude |
| 4 | FE_UR | -1.628 | -1.454 | -1.067 | -2.639 | -1.746 | 0.679 | 0.321 | coex | 0.873 | 0.312 | coex |
| 5 | FE_UR | -1.914 | -1.498 | -1.562 | -1.480 | -0.335 | -0.052 | 1.052 | exclude | 0.168 | -0.185 | exclude |
| 6 | FE_UR | -2.499 | -1.247 | -1.431 | -2.162 | -1.984 | 0.461 | 0.539 | coex | 0.992 | -0.077 | coex |
| 7 | FE_UR | -1.901 | -0.952 | -1.730 | -1.314 | -0.532 | 0.680 | 0.320 | coex | 0.266 | 0.096 | coex |
| 8 | FE_UR | -1.637 | -1.642 | -1.079 | -0.956 | 0.128 | 5.352 | -4.352 | exclude | -0.064 | -0.622 | exclude |
| 9 | FE_UR | -1.146 | -0.341 | -2.276 | -1.441 | 0.031 | -35.453 | 36.453 | exclude | -0.016 | 1.115 | exclude |
| 3 | HO_PL | -1.463 | -0.965 | -1.564 | -1.486 | -0.420 | 1.241 | -0.241 | exclude | 0.210 | 0.311 | exclude |
| 4 | HO_PL | -0.843 | -0.659 | -1.896 | -2.242 | -0.530 | 2.984 | -1.984 | exclude | 0.265 | 1.318 | exclude |
| 5 | HO_PL | -1.570 | -1.472 | -1.398 | -1.570 | -0.271 | 0.362 | 0.638 | coex | 0.135 | -0.037 | coex |
| 6 | HO_PL | -1.138 | -1.108 | -1.699 | -1.843 | -0.174 | 4.229 | -3.229 | exclude | 0.087 | 0.648 | exclude |
| 7 | HO_PL | -0.955 | -1.201 | -2.395 | -2.069 | 0.572 | -1.516 | 2.516 | exclude | -0.286 | 1.154 | exclude |
| 8 | HO_PL | -0.914 | -0.959 | -1.922 | -2.291 | -0.324 | 4.110 | -3.110 | exclude | 0.162 | 1.170 | exclude |
| 9 | HO_PL | -1.192 | -0.864 | -1.255 | -2.326 | -1.399 | 1.045 | -0.045 | exclude | 0.700 | 0.762 | exclude |
| 10 | HO_PL | -1.101 | -1.034 | -1.462 | -2.485 | -1.090 | 1.331 | -0.331 | exclude | 0.545 | 0.905 | exclude |
| 1 | HO_SA | -0.887 | -0.500 | -1.406 | -0.915 | 0.103 | -4.043 | 5.043 | exclude | -0.051 | 0.467 | exclude |
| 3 | HO_SA | -1.463 | -0.940 | -1.540 | -1.961 | -0.944 | 1.082 | -0.082 | exclude | 0.472 | 0.549 | exclude |
| 4 | HO_SA | -0.843 | -0.538 | -1.590 | -1.112 | 0.172 | -3.340 | 4.340 | exclude | -0.086 | 0.661 | exclude |
| 5 | HO_SA | -1.570 | -1.115 | -1.656 | -1.421 | -0.220 | 1.391 | -0.391 | exclude | 0.110 | 0.196 | exclude |
| 7 | HO_SA | -0.955 | -1.076 | -1.809 | -1.624 | 0.306 | -1.788 | 2.788 | exclude | -0.153 | 0.701 | exclude |
| 8 | HO_SA | -0.914 | -0.651 | -1.977 | -2.156 | -0.442 | 3.404 | -2.404 | exclude | 0.221 | 1.285 | exclude |
| 9 | HO_SA | -1.192 | -0.513 | -1.679 | -1.439 | -0.439 | 2.110 | -1.110 | exclude | 0.219 | 0.706 | exclude |
| 10 | HO_SA | -1.101 | -1.138 | -1.770 | -1.144 | 0.662 | -0.009 | 1.009 | exclude | -0.331 | 0.337 | exclude |
| 1 | HO_UR | -0.887 | -0.551 | -2.919 | -1.695 | 0.888 | -1.289 | 2.289 | exclude | -0.444 | 1.588 | exclude |
| 2 | HO_UR | -1.088 | -1.016 | -0.910 | -1.189 | -0.351 | 0.490 | 0.510 | coex | 0.175 | -0.003 | coex |
| 4 | HO_UR | -0.843 | -0.666 | -1.634 | -2.639 | -1.182 | 1.669 | -0.669 | exclude | 0.591 | 1.382 | exclude |
| 5 | HO_UR | -1.570 | -1.118 | -1.212 | -1.480 | -0.720 | 0.503 | 0.497 | coex | 0.360 | 0.002 | coex |
| 6 | HO_UR | -1.138 | -0.903 | -1.735 | -2.162 | -0.663 | 1.900 | -0.900 | exclude | 0.331 | 0.928 | exclude |
| 7 | HO_UR | -0.955 | -0.824 | -1.068 | -1.314 | -0.377 | 1.299 | -0.299 | exclude | 0.188 | 0.301 | exclude |

| | | | | | | | | | | | | |
|---|---|---|---|---|---|---|---|---|---|---|---|---|
| 8 | HO_UR | -0.914 | -0.450 | -1.130 | -0.956 | -0.290 | 1.746 | -0.746 | exclude | 0.145 | 0.361 | exclude |
| 9 | HO_UR | -1.192 | -0.660 | -1.123 | -1.441 | -0.850 | 0.919 | 0.081 | coex | 0.425 | 0.356 | coex |
| 1 | PL_SA | -1.422 | -1.175 | -1.914 | -0.915 | 0.752 | 0.346 | 0.654 | exclude | -0.376 | 0.116 | exclude |
| 3 | PL_SA | -1.486 | -1.721 | -2.055 | -1.961 | 0.329 | -0.730 | 1.730 | exclude | -0.164 | 0.404 | exclude |
| 4 | PL_SA | -2.242 | -1.653 | -1.497 | -1.112 | -0.204 | -2.653 | 3.653 | exclude | 0.102 | -0.643 | exclude |
| 5 | PL_SA | -1.570 | -1.229 | -1.692 | -1.421 | -0.070 | 2.742 | -1.742 | exclude | 0.035 | 0.157 | exclude |
| 7 | PL_SA | -2.069 | -1.878 | -1.680 | -1.624 | -0.135 | -1.873 | 2.873 | exclude | 0.068 | -0.322 | exclude |
| 8 | PL_SA | -2.291 | -0.963 | -1.977 | -2.156 | -1.507 | 0.792 | 0.208 | coex | 0.754 | 0.440 | coex |
| 9 | PL_SA | -2.326 | -0.882 | -2.238 | -1.439 | -0.645 | 0.863 | 0.137 | coex | 0.322 | 0.234 | coex |
| 10 | PL_SA | -2.485 | -1.749 | -1.925 | -1.144 | 0.045 | 13.419 | -12.419 | exclude | -0.023 | -0.582 | exclude |
| 1 | PL_UR | -1.422 | -1.637 | -2.164 | -1.695 | 0.684 | -0.085 | 1.085 | exclude | -0.342 | 0.400 | exclude |
| 2 | PL_UR | 0.324 | -1.311 | -1.309 | -1.189 | 1.756 | 0.070 | 0.930 | exclude | -0.878 | 0.755 | exclude |
| 4 | PL_UR | -2.242 | -2.046 | -1.677 | -2.639 | -1.158 | 0.512 | 0.488 | coex | 0.579 | 0.014 | coex |
| 5 | PL_UR | -1.570 | -0.674 | -1.518 | -1.480 | -0.858 | 0.939 | 0.061 | coex | 0.429 | 0.377 | coex |
| 6 | PL_UR | -1.843 | -1.516 | -1.794 | -2.162 | -0.695 | 0.929 | 0.071 | coex | 0.348 | 0.299 | coex |
| 7 | PL_UR | -2.069 | -1.125 | -1.592 | -1.314 | -0.666 | 0.284 | 0.716 | coex | 0.333 | -0.144 | coex |
| 8 | PL_UR | -2.291 | -1.333 | -2.371 | -0.956 | 0.457 | 0.825 | 0.175 | exclude | -0.229 | -0.148 | exclude |
| 9 | PL_UR | -2.326 | -1.316 | -1.863 | -1.441 | -0.588 | 0.213 | 0.787 | coex | 0.294 | -0.169 | coex |
| 1 | SA_UR | -0.915 | -0.645 | -1.493 | -1.695 | -0.473 | 2.220 | -1.220 | exclude | 0.237 | 0.814 | exclude |
| 4 | SA_UR | -1.112 | -0.644 | -1.304 | -2.639 | -1.803 | 1.106 | -0.106 | exclude | 0.902 | 1.093 | exclude |
| 5 | SA_UR | -1.421 | -1.394 | -1.188 | -1.480 | -0.320 | 0.270 | 0.730 | coex | 0.160 | -0.073 | coex |
| 7 | SA_UR | -1.624 | -1.709 | -1.217 | -1.314 | -0.012 | -32.707 | 33.707 | exclude | 0.006 | -0.401 | exclude |
| 8 | SA_UR | -2.156 | -1.223 | -0.755 | -0.956 | -1.135 | -0.235 | 1.235 | exclude | 0.567 | -0.834 | exclude |
| 9 | SA_UR | -1.439 | -1.671 | -1.814 | -1.441 | 0.604 | 0.380 | 0.620 | exclude | -0.302 | 0.073 | exclude |

### Appendix S4: R Code to recreate analysis from this paper

```
knitr::opts_chunk$set(warning = FALSE)
# Load libraries ------
library(tidyverse)
library(patchwork)
# Theme for plots
theme_gsk <- function() {
  theme_minimal()+
    theme(panel.border = element_blank(),
          panel.grid.major = element_blank(),
          panel.grid.minor = element_blank(),
          axis.line = element_line(colour = "black"),
          plot.tag = element_text(face = "bold")
    )
}

# Toggle to true if you want script to generate files/figures
# leave it on false if you don't want to override existing files.
write_objects <- FALSE
```

We are now ready to read in data:

```
# Read in the data and clean out empty rows. ------
biomass <- read_csv("../../data/phase2-harvest.csv")
```

```
## Parsed with column specification:
## cols(
##   number = col_double(),
##   pot_id = col_character(),
##   source_soil = col_character(),
##   focal_species = col_character(),
##   replicate = col_character(),
##   abg_dry_g = col_double()
## )
```

```
# Get rid of "empty" rows
biomass <- biomass %>% arrange(focal_species, source_soil, replicate) %>%
  filter(!(grepl("Empty", biomass$source_soil)))
# Get a sense for the structure of the dataframe and the data itself

glimpse(biomass)
```

```
## Observations: 480
## Variables: 6
## $ number        <dbl> 39, 471, 53, 129, 182, 233, 271, 324, 383, 425, 12, 4...
## $ pot_id        <chr> "aastr_ACWR_R1", "aastr_ACWR_R10", "aastr_ACWR_R2", "...
## $ source_soil   <chr> "aastr", "aastr", "aastr", "aastr", "aastr", "aastr",...
## $ focal_species <chr> "ACWR", "ACWR", "ACWR", "ACWR", "ACWR", "ACWR", "ACWR...
## $ replicate     <chr> "R1", "R10", "R2", "R3", "R4", "R5", "R6", "R7", "R8"...
## $ abg_dry_g     <dbl> 0.0407, 0.0240, 0.0363, 0.0429, 0.0368, 0.0388, 0.027...
```

```
# 6 columns:
# 1. "number" - this just has a unique number for each pot, and was used to randomize the map
# 1. cont. Not really meaningful for this analysis, except as a way to keep track of each pot.
# 2. pot_id - the unique ID for each pot. soil_focalspecies_replicate
```

```r
# 3. Source soil - identifies which soil was used to inoculate the pot
# 3. cont. Eight unique soils: AC, FE, HO, SA, PL, UR, and two controls.
# 3. cont. the two controls are "aastr" and "abfld" and refer to sterile and field soil.
# 4. focal_species: identity of the species growing in the pot.
# 5. replicate: which block the pot came from.
# 6. abg_dry_g: dry biomass, in grams.

# There are 48 combinations of soil x focal species, grown in ten replicate racks
# for a total of 480 rows:
dim(biomass)
```

```
## [1] 480   6
```

```r
# Unfortunately 6 pots did not have live plants at the two-week mark,
# so these plots will be excluded from further analysis:
biomass %>% filter(is.na(abg_dry_g)) %>% select(pot_id)
```

```
## # A tibble: 6 x 1
##   pot_id
##   <chr>
## 1 PL_HOMU_R1
## 2 HO_PLER_R2
## 3 abfld_SACO_R2
## 4 abfld_SACO_R6
## 5 abfld_URLI_R10
## 6 UR_URLI_R3
```

```r
num_NA_pots_main <- biomass %>% filter(is.na(abg_dry_g)) %>% nrow
# We can eliminate these pots from the datafame:

biomass <- na.omit(biomass)

# Let's explore the distribution of the data. ------
# Exploring the distribution of biomasses
biom_hist <- ggplot(biomass) +
  geom_histogram(aes(x = abg_dry_g)) +
  facet_wrap(.~focal_species)
biom_hist_log <- biom_hist + scale_x_log10() + ggtitle("logged x-axis")

biom_hist/biom_hist_log
```

```
## `stat_bin()` using `bins = 30`. Pick better value with `binwidth`.

## `stat_bin()` using `bins = 30`. Pick better value with `binwidth`.
```

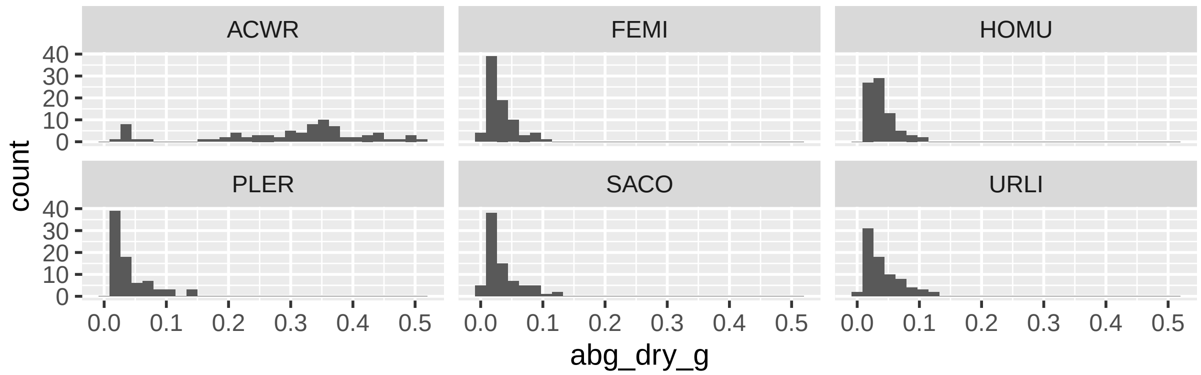

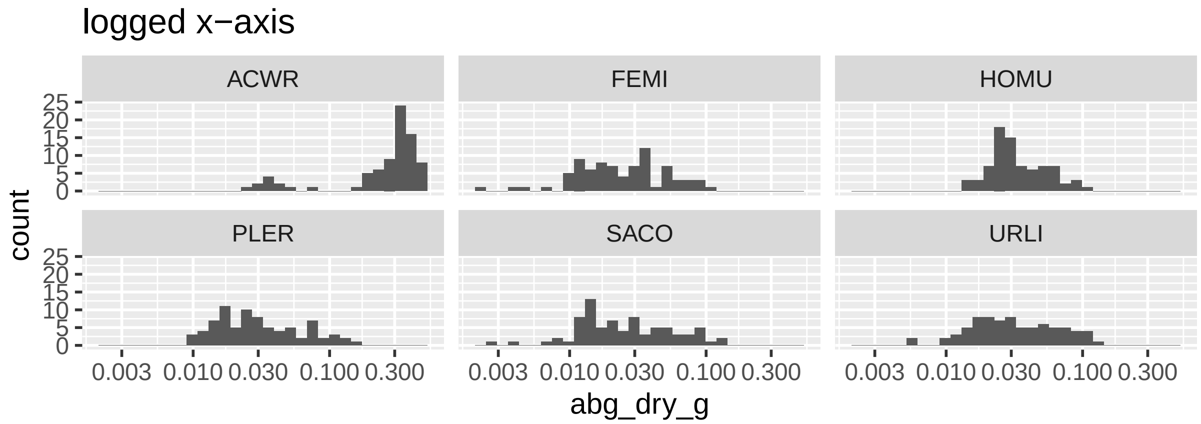

```
# Rename columns with more intuitive names - helps down the line
biomass <- biomass %>%
  # Rename soil names; add a column that adds consp or hetsp source
  mutate(pointcol = ifelse(source_soil == "aastr", "Sterile",
                           "Phase 1\nHeterospecific\ncultivated"),
         pointcol = ifelse(source_soil == "abfld", "Field", pointcol),
         pointcol = ifelse(str_extract(focal_species, "..") == source_soil,
                           "Phase 1\nConspecific\ncultivated", pointcol),
         source_soil = ifelse(source_soil == "abfld", "Field", source_soil),
         source_soil = ifelse(source_soil == "aastr", "Sterile", source_soil))
```

Now that the data are imported and reshaped in a workable way, we can plot the biomasses of each plant species in each soil type (Figure 1 of manuscript):

```
# Make a "Cleveland dotplot" type plot of the biomass values (Figure 1) -------
# Point will be plotted at the Median, with error bars extending
# from the LQR to the UQR.
# When maximum or minimum values in a group are more then 1.5*IQR
# away from L/UQR, they will be plotted.
biomass_for_cleveland <- biomass %>%
  group_by(focal_species, source_soil, pointcol) %>%
  summarize(
    # Get the mean and SEM for each (focal*soil source) group:
    mean_bm = mean(abg_dry_g),
    se_bm = sd(abg_dry_g)/sqrt(n()),
    # Also get the median and IQRs
    median_bm = median(abg_dry_g),
    lqr = quantile(abg_dry_g, 0.25),
    uqr = quantile(abg_dry_g, 0.75),
```

```
    # Get outliers
    min_val = min(abg_dry_g, na.rm = T),
    max_val = max(abg_dry_g, na.rm = T),
    out_low = ifelse(min_val < lqr - 1.5*(uqr-lqr), min_val, NA),
    out_upp = ifelse(max_val > uqr + 1.5*(uqr-lqr), max_val, NA),
    n = n())

# add a column that sets the y-value of the points --
# needed, because I want to add the outlier points
biomass_for_cleveland$plot_y <- rev(c(seq(0.6, 1.3, by = .1),
                                      seq(1.6, 2.3, by = .1)+1,
                                      seq(2.6, 3.3, by = .1)+2,
                                      seq(3.6, 4.3, by = .1)+3,
                                      seq(4.6, 5.3, by = .1)+4,
                                      seq(5.6, 6.3, by = .1)+5))
# Make a new data frame that includes the outlier points
# Outliers here are defined as those points that are either
# lower than (LQR - 1.5*IQR), or higher than (UQR + 1.5*IQR)
outliers <- left_join(biomass, biomass_for_cleveland) %>%
  filter(abg_dry_g < lqr - (1.5*(uqr-lqr)) |
           abg_dry_g > uqr + (1.5*(uqr-lqr))) %>%
  select(focal_species, source_soil, abg_dry_g, plot_y)
```

```
## Joining, by = c("source_soil", "focal_species", "pointcol")
```

```
(biomass_dotplot <- ggplot(biomass_for_cleveland,
                          aes(y = plot_y, x = median_bm, fill = pointcol,
                              label = source_soil, shape = pointcol)) +
    scale_x_log10() +
    # add median value
    geom_point(size = 4, stroke = .9) +
    # add dashed lines connecting outlier to median (if outlier exists)
    geom_errorbarh(aes(xmin = out_low, xmax = lqr),
                   height = 0, linetype = 3, size = .25) +
    geom_errorbarh(aes(xmin = uqr, xmax = out_upp),
                   height = 0, linetype = 3, size = .25) +
    # add outlying points (if they exist)
    geom_point(aes(y = plot_y, x = out_low), shape = 21, fill = "grey50", size = .5) +
    geom_point(aes(y = plot_y, x = out_upp), shape = 21, fill = "grey50", size = .5) +
    geom_point(data = outliers, aes(x = abg_dry_g, y = plot_y),
               shape = 21, fill = "grey50", size = .5, inherit.aes = F) +

    # add solid line connecting median to lower and upper quantile
    geom_errorbarh(aes(xmin = lqr, xmax = uqr), height = .1) +

    # Set color and shape for all the points
    scale_fill_manual(values = c("#009E73", "grey10", "grey90", "#D55E00"),
                      name = "Inoculum\nsource") +
    scale_shape_manual(values = c(22,21,21,23),
                       name = "Inoculum\nsource") +
    # add thin line between each focal species
    geom_hline(yintercept = c(2, 4, 6, 8, 10), linetype = 1, size = 0.25) +
```

```
  # add species names along y-axis
  scale_y_continuous(breaks = c(1,3,5,7,9,11),
                     labels = rev(unique(biomass_for_cleveland$focal_species))) +
  # misc. theme settings.
  ylab("Focal species") +
  xlab("Aboveground biomass (g)") +
  theme_gsk() +
  theme(#legend.justification=c(1,0), legend.position=c(1,.15),
    legend.position = "top",
    # legend.background = element_rect(colour = "grey50"),
    legend.text = element_text(size = 8)) +
  NULL)
```

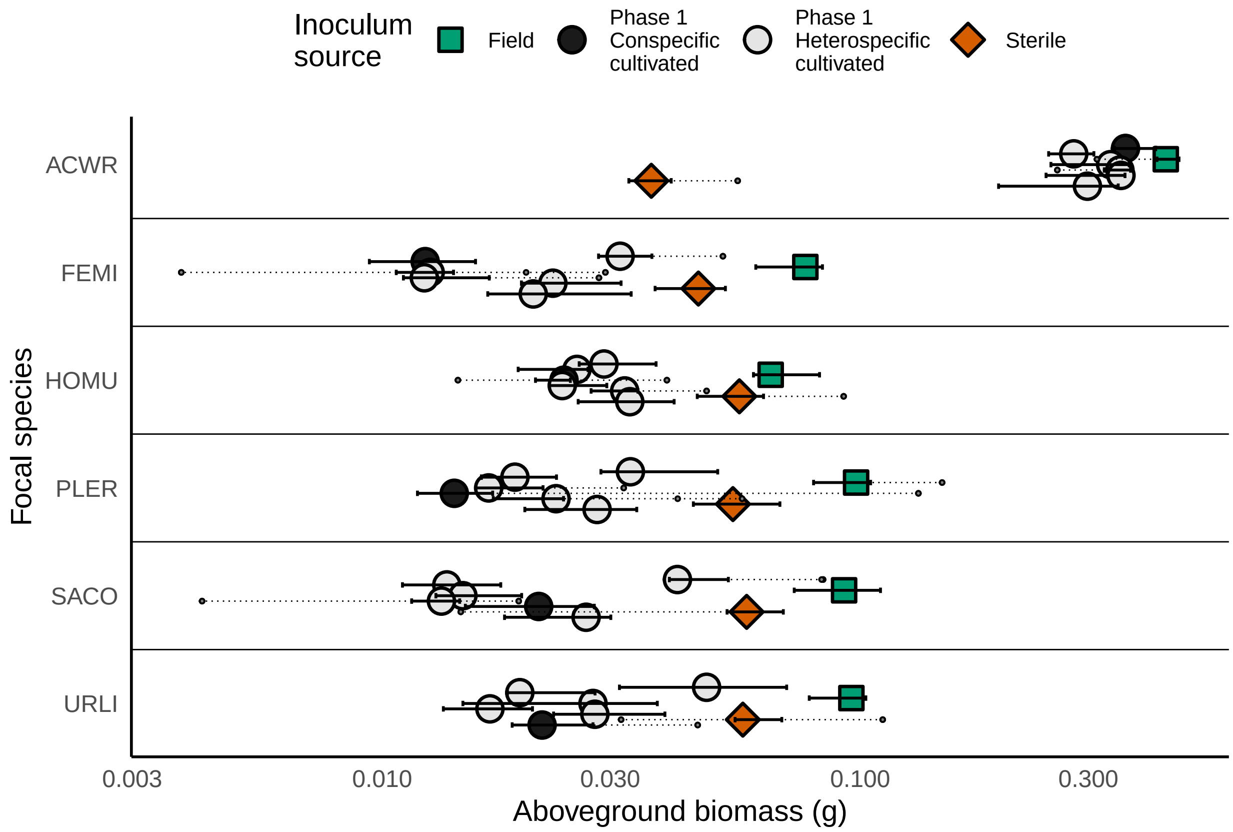

```
if(write_objects) {
  saveRDS(biomass_dotplot, file = "../../figures/biomass_dotplot.Rds")
}
```

The main goal now is to go from these biomass values to estimtates of the key $m$ parameters – and from there, to quantify the degree of microbially mediated stabilization and fitness differences.

```
# Now, the goal is to use the biomass values to estimate
# the degree of microbially mediated stabilization and
# fitness differences.
# Reshaping data for calculating Stabilization and FDs ----

# So first, let's make a column that is simply the log AGB
# of each pot, and then go from there.
biomass$log_agb <- log(biomass$abg_dry_g)
```

```r
# Now, we need to convert the "long format" biomass data frame
# into a "wide format". Here's a function for that:
make_wide_biomass <- function(df) {
  # This takes the biomass dataframe, and converts into a wide
  # dataframe, such that each row is one replicate rack,
  # and each column represents the log AGB of one species in
  # one soil background in that rack.
  df %>% mutate(source_soil = ifelse(source_soil == "aastr", "str", source_soil),
                source_soil = ifelse(source_soil == "abfld", "fld", source_soil),
                pair = paste0(source_soil, "_", focal_species)) %>%
    select(replicate, pair, log_agb) %>%
    spread(pair, log_agb)
}
biomass_wide <- make_wide_biomass(biomass)

# Calculate the Stabilization between each species pair ----
# Recall that following the definition of the m terms in Bever 1997,
# and the analysis of this model in Kandlikar 2019,
# stablization = -0.5*(log(m1A) + log(m2B) - log(m1B) - log(m2A))
# Here is a function that does this calculation
# (recall that after the data reshaping above,
# each column represents a given value of log(m1A))
calculate_stabilization <- function(df) {
  df %>%
    # In df, each row is one repilcate/rack, and each column
    # represents the growth of one species in one soil type.
    # e.g. "FE_ACWR" is the growth of FE in ACWR-cultivated soil.
    mutate(AC_FE = -0.5*(AC_ACWR - AC_FEMI - FE_ACWR + FE_FEMI),
           AC_HO = -0.5*(AC_ACWR - AC_HOMU - HO_ACWR + HO_HOMU),
           AC_SA = -0.5*(AC_ACWR - AC_SACO - SA_ACWR + SA_SACO),
           AC_PL = -0.5*(AC_ACWR - AC_PLER - PL_ACWR + PL_PLER),
           AC_UR = -0.5*(AC_ACWR - AC_URLI - UR_ACWR + UR_URLI),
           FE_HO = -0.5*(FE_FEMI - FE_HOMU - HO_FEMI + HO_HOMU),
           FE_SA = -0.5*(FE_FEMI - FE_SACO - SA_FEMI + SA_SACO),
           FE_PL = -0.5*(FE_FEMI - FE_PLER - PL_FEMI + PL_PLER),
           FE_UR = -0.5*(FE_FEMI - FE_URLI - UR_FEMI + UR_URLI),
           HO_PL = -0.5*(HO_HOMU - HO_PLER - PL_HOMU + PL_PLER),
           HO_SA = -0.5*(HO_HOMU - HO_SACO - SA_HOMU + SA_SACO),
           HO_UR = -0.5*(HO_HOMU - HO_URLI - UR_HOMU + UR_URLI),
           SA_PL = -0.5*(SA_SACO - SA_PLER - PL_SACO + PL_PLER),
           SA_UR = -0.5*(SA_SACO - SA_URLI - UR_SACO + UR_URLI),
           PL_UR = -0.5*(PL_PLER - PL_URLI - UR_PLER + UR_URLI)) %>%
    select(replicate, AC_FE:PL_UR) %>%
    gather(pair, stabilization, AC_FE:PL_UR)
}

stabilization_values <- calculate_stabilization(biomass_wide)
# Seven values are NA; we can omit these
stabilization_values <- stabilization_values %>%
  filter(!(is.na(stabilization)))

# Calculating the Fitness difference between each species pair ----
# Similarly, we can now calculate the fitness difference between
```

```r
# each pair. Recall that FD = 0.5*(log(m1A)+log(m1B)-log(m2A)-log(m2B))
# But recall that here, IT IS IMPORTANT THAT
# m1A = (m1_soilA - m1_fieldSoil)!

# The following function does this calculation:
calculate_fitdiffs <- function(df) {
  df %>%
    mutate(AC_FE = 0.5*((AC_ACWR-Field_ACWR) + (FE_ACWR-Field_ACWR) -
                      (AC_FEMI-Field_FEMI) - (FE_FEMI-Field_FEMI)),
           AC_HO = 0.5*((AC_ACWR-Field_ACWR) + (HO_ACWR-Field_ACWR) -
                      (AC_HOMU-Field_HOMU) - (HO_HOMU-Field_HOMU)),
           AC_SA = 0.5*((AC_ACWR-Field_ACWR) + (SA_ACWR-Field_ACWR) -
                      (AC_SACO-Field_SACO) - (SA_SACO-Field_SACO)),
           AC_PL = 0.5*((AC_ACWR-Field_ACWR) + (PL_ACWR-Field_ACWR) -
                      (AC_PLER-Field_PLER) - (PL_PLER-Field_PLER)),
           AC_UR = 0.5*((AC_ACWR-Field_ACWR) + (UR_ACWR-Field_ACWR) -
                      (AC_URLI-Field_URLI) - (UR_URLI-Field_URLI)),
           FE_HO = 0.5*((FE_FEMI-Field_FEMI) + (HO_FEMI-Field_FEMI) -
                      (FE_HOMU-Field_HOMU) - (HO_HOMU-Field_HOMU)),
           FE_SA = 0.5*((FE_FEMI-Field_FEMI) + (SA_FEMI-Field_FEMI) -
                      (FE_SACO-Field_SACO) - (SA_SACO-Field_SACO)),
           FE_PL = 0.5*((FE_FEMI-Field_FEMI) + (PL_FEMI-Field_FEMI) -
                      (FE_PLER-Field_PLER) - (PL_PLER-Field_PLER)),
           FE_UR = 0.5*((FE_FEMI-Field_FEMI) + (UR_FEMI-Field_FEMI) -
                      (FE_URLI-Field_URLI) - (UR_URLI-Field_URLI)),
           HO_PL = 0.5*((HO_HOMU-Field_HOMU) + (PL_HOMU-Field_HOMU) -
                      (HO_PLER-Field_PLER) - (PL_PLER-Field_PLER)),
           HO_SA = 0.5*((HO_HOMU-Field_HOMU) + (SA_HOMU-Field_HOMU) -
                      (HO_SACO-Field_SACO) - (SA_SACO-Field_SACO)),
           HO_UR = 0.5*((HO_HOMU-Field_HOMU) + (UR_HOMU-Field_HOMU) -
                      (HO_URLI-Field_URLI) - (UR_URLI-Field_URLI)),
           SA_PL = 0.5*((SA_SACO-Field_SACO) + (PL_SACO-Field_SACO) -
                      (SA_PLER-Field_PLER) - (PL_PLER-Field_PLER)),
           SA_UR = 0.5*((SA_SACO-Field_SACO) + (UR_SACO-Field_SACO) -
                      (SA_URLI-Field_URLI) - (UR_URLI-Field_URLI)),
           PL_UR = 0.5*((PL_PLER-Field_PLER) + (UR_PLER-Field_PLER) -
                      (PL_URLI-Field_URLI) - (UR_URLI-Field_URLI))) %>%
    select(replicate, AC_FE:PL_UR) %>%
    gather(pair, fitdiff_fld, AC_FE:PL_UR)
}
fd_values <- calculate_fitdiffs(biomass_wide)
# Twenty-two values are NA; let's omit these.
fd_values <- fd_values %>% filter(!(is.na(fitdiff_fld)))

# Now, generate statistical summaries of SD and FD

stabiliation_summary <- stabilization_values %>% group_by(pair) %>%
  summarize(mean_sd = mean(stabilization),
            sem_sd = sd(stabilization)/sqrt(n()),
            n_sd = n())

fitdiff_summary <- fd_values %>% group_by(pair) %>%
  summarize(mean_fd = mean(fitdiff_fld),
```

```
            sem_fd = sd(fitdiff_fld)/sqrt(n()),
            n_fd = n())

# Combine the two separate data frames.
sd_fd_summary <- left_join(stabiliation_summary, fitdiff_summary)
```

```
## Joining, by = "pair"
```

```
# Some of the FDs are negative, let's flip these to be positive
# and also flip the label so that the first species in the name
# is always the fitness superior.
sd_fd_summary <- sd_fd_summary %>%
  # if mean_fd is < 0, the following command gets the absolute
  # value and also flips around the species code so that
  # the fitness superior is always the first species in the code
  mutate(pair = ifelse(mean_fd < 0,
                       paste0(str_extract(pair, "..$"),
                              "_",
                              str_extract(pair, "^..")),
                       pair),
         mean_fd = abs(mean_fd))

# Make a column that indicates the net outcome (coex. or exclusion)
# to do so, check if the stabilization estimate is larger
# or smaller than the FD estimate.
sd_fd_summary <- sd_fd_summary %>% mutate(
  stabilize = ifelse(mean_sd - 2*sem_sd > 0, "yes", "no"),
  fitness = ifelse(mean_fd - 2*sem_fd > 0, "yes", "no"),
  outcome = ifelse(mean_fd - 2*sem_fd >
                     mean_sd + 2*sem_sd, "exclusion", "neutral"),
  outcome2 = ifelse(mean_fd > mean_sd, "exclusion", "coexistence"))
head(sd_fd_summary)
```

```
## # A tibble: 6 x 11
##   pair  mean_sd sem_sd  n_sd mean_fd sem_fd  n_fd stabilize fitness outcome
##   <chr>   <dbl>  <dbl> <int>   <dbl>  <dbl> <int> <chr>     <chr>   <chr>
## 1 AC_FE  0.376  0.0864    10   1.01  0.112     10 yes       yes     exclus~
## 2 AC_HO  0.0455 0.0490    10   0.724 0.0765    10 no        yes     exclus~
## 3 AC_PL  0.341  0.132     10   1.14  0.148     10 yes       yes     exclus~
## 4 AC_SA  0.333  0.0825    10   0.789 0.213      8 yes       yes     neutral
## 5 AC_UR  0.227  0.148      9   0.834 0.183      8 no        yes     neutral
## 6 HO_FE  0.0311 0.0731    10   0.689 0.147     10 no        yes     exclus~
## # ... with 1 more variable: outcome2 <chr>
```

We are now ready to plot the strength of microbially mediated stabilization against the strength of microbially mediated fitness differences.

```
# Plot SD vs. FD -----------
(sd_fd_xpyplot <- ggplot(sd_fd_summary, aes(x = mean_sd, y = mean_fd, label = pair,
                                            fill = outcome)) +
  geom_ribbon(aes(x = seq(0, 1.55, length.out = 15),
                  ymin = rep(0, 15),
                  ymax = seq(0, 1.55, length.out = 15)),
              fill = "#FFF2CC")  +
  geom_errorbar(aes(ymin = (mean_fd)-2*sem_fd,
```

```
                  ymax = (mean_fd)+2*sem_fd),
              size = .25) +
geom_errorbarh(aes(xmin = mean_sd-2*sem_sd,
                   xmax = mean_sd+2*sem_sd),
               size = .25) +
ggrepel::geom_text_repel(segment.size = 0.025, color = "#610B0B", size = 3,
                         nudge_x = c(.15, -.2, .12, .2, .3,
                                     -0.2, -.25, -.05, .28, -.1,
                                     -.2, 0.15, .20, -.15, -.15),
                         nudge_y = c(.1, .08, .12, -.05, .05,
                                     -.075, -.45, .1, .1, .2,
                                     -.05, -.05, -.25, .08, 0.1)
) +
xlim(c(-.5, 1.25)) +
geom_abline(linetype = 2) +
# geom_label() +
geom_hline(yintercept = 0) + geom_vline(xintercept = 0)+
geom_point(shape = 21, size = 2, stroke = 1) +
scale_fill_manual(values = c("grey50", "white"),
                  labels = c("PSFs drive exclusion",
                             "No strong evidence that PSFs\ndrive exclusion or coexistence"),
                  name = "Coexistence outcome") +
xlab(bquote(atop("Microbially mediated stabilization",
                 -frac(1,2)~(m["1A"]-m["1B"]-m["2A"]+m["2B"]) ))) +
ylab(bquote(atop("Microbially mediated fitness difference",
                 frac(1,2)~(m["1A"]+m["1B"]-m["2A"]-m["2B"])))) +
annotate("text", x = 1.1, y = .05, label = "coexistence",
         vjust = 0, hjust = 1, size = 4, fontface = "bold.italic") +
annotate("text", x = 1.1, y = 1.25, label = "exclusion",
         vjust = 0, hjust = 1, size = 4, fontface = "bold.italic") +

theme_gsk() +
theme(axis.line = element_line(size = 0),
      legend.justification=c(1,0), legend.position=c(1,.025),
      legend.background = element_rect(colour = "grey50")) +
NULL)
```

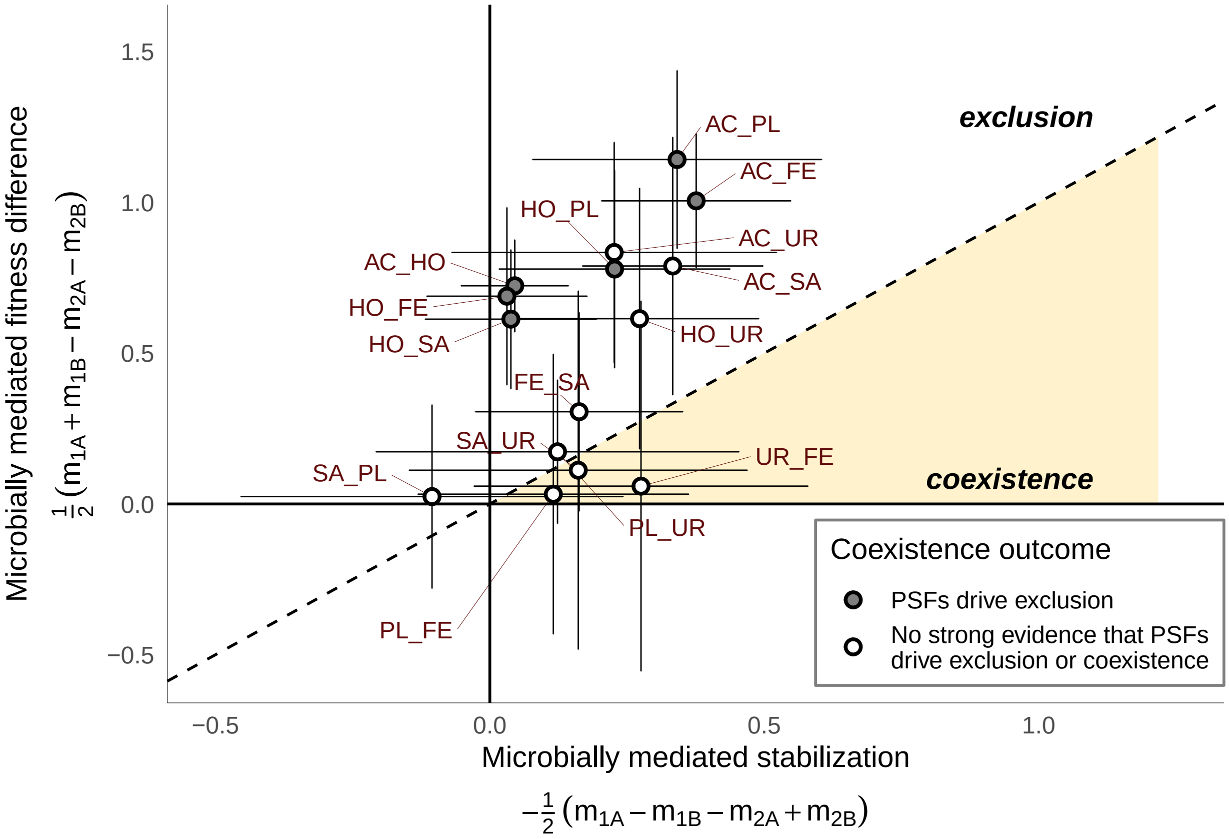

```
if(write_objects) {
  saveRDS(sd_fd_xpyplot, file = "../figures/sd_fd_xyplot.Rds")
}
```

Next, we can explore the dataset in a few ways that help us build a meaningful discussion:

```
# Now, answering some basic questions: ------
# 1. How much lower was the biomass of ACWR in sterile soil than
# average growth with any live soil?
acwr_sterile_mean <- biomass %>% filter(focal_species == "ACWR" &
                     source_soil == "Sterile") %>%
   summarize(mean = mean(abg_dry_g)) %>% unlist

acwr_live_mean <- biomass %>% filter(focal_species == "ACWR" &
                     source_soil != "Sterile") %>%
  summarize(mean = mean(abg_dry_g)) %>% unlist

acwr_field_mean <- biomass %>% filter(focal_species == "ACWR" &
                                  source_soil == "Field") %>%
  summarize(mean = mean(abg_dry_g)) %>% unlist
acwr_nonFieldLive_mean <- biomass %>% filter(focal_species == "ACWR" &
                                        source_soil != "Field" &
                                         source_soil != "Sterile") %>%
  summarize(mean = mean(abg_dry_g)) %>% unlist
acwr_field_nonField_ratio <- acwr_nonFieldLive_mean/acwr_field_mean

nonacwr_field_mean <- biomass %>% filter(focal_species != "ACWR" &
```

```
                                       source_soil == "Field") %>%
  summarize(mean = mean(abg_dry_g)) %>% unlist
nonacwr_nonFieldLive_mean <- biomass %>% filter(focal_species != "ACWR" &
                                              source_soil != "Field" &
                                              source_soil != "Sterile") %>%
  summarize(mean = mean(abg_dry_g)) %>% unlist
nonacwr_field_nonField_ratio <- nonacwr_nonFieldLive_mean/nonacwr_field_mean

# 2. How different is growth in conspecific soils
# vs. growth in heterospecific soils, across all species?
mean_growth_in_conspecific <- biomass %>%
  filter(pointcol == "Phase 1\nConspecific\ncultivated") %>%
  group_by(focal_species) %>% summarize(mean_self = mean(abg_dry_g)) %>%
  select(mean_self) %>% unlist
mean_growth_in_heterospecific <- biomass %>%
  filter(pointcol == "Phase 1\nHeterospecific\ncultivated") %>%
  group_by(focal_species) %>% summarize(mean_nonself = mean(abg_dry_g)) %>%
  select(mean_nonself) %>% unlist

# 3. What is the outcome of running an ANOVA
# for the main experiment?
biomass_lm <- lm(log(abg_dry_g) ~ source_soil*focal_species, biomass)
biomass_lm_anova <- car::Anova(biomass_lm)

# Save these one-off objects as a list to be called in the
# RMarkdown file.
if(write_objects) {
  saveRDS(object = list(biomass_lm_anova = biomass_lm_anova,
                        nonacwr_field_nonField_ratio = nonacwr_field_nonField_ratio,
                        acwr_field_nonField_ratio = acwr_field_nonField_ratio,
                        mean_growth_in_conspecific = mean_growth_in_conspecific,
                        mean_growth_in_heterospecific = mean_growth_in_heterospecific),
          file = "../figures/relevant_objects.Rds")

}
```

We can also make a table that displays the mean and confidence interval of each FD and stabilization estimate.

```
# Make a table of outcomes (table 1 of paper) --------
tbl_to_print <- sd_fd_summary %>%
  mutate(`Species pair` = c("\\textit{A. wrangelianus/\nF.  microstachys}",
                            "\\textit{A. wrangelianus/\nH.  murinum}",
                            "\\textit{A. wrangelianus/\nP.  erecta}",
                            "\\textit{A. wrangelianus/\nS. columbariae}",
                            "\\textit{A. wrangelianus/\nU.  lindleyi}",
                            "\\textit{H.  murinum/\nF.  microstachys}",
                            "\\textit{P.  erecta/\nF.  microstachys}",
                            "\\textit{F.  microstachys/\nS. columbariae}",
                            "\\textit{U.  lindleyi/\nF.  microstachys}",
                            "\\textit{H.  murinum/\nP.  erecta}",
                            "\\textit{H.  murinum/\nS. columbariae}",
                            "\\textit{H.  murinum/\nU.  lindleyi}",
                            "\\textit{P.  erecta/\nU.  lindleyi}",
```

```
                              "\\textit{S. columbariae/\nP.  erecta}",
                              "\\textit{S. columbariae/\nU.  lindleyi}")) %>%
  mutate_if(is.numeric, round, 3) %>%
  mutate(arrangement = c(3,3,3, 2,2,3, 1,2,1, 3,3,2, 2,2,1)) %>%
  arrange(arrangement) %>%
  mutate(pair = str_replace(pair, "_", "\\\\_")) %>%
  mutate(outcome2 = ifelse(outcome == "exclusion", paste0("\\textbf{", outcome2, "}"), outcome2)) %>%
  mutate(Stabilization = ifelse(mean_sd - 2*sem_sd > 0, paste0("\\textbf{",mean_sd,"}"), mean_sd)) %>%
  mutate(`Fitness Difference` = ifelse(mean_fd - 2*sem_fd > 0, paste0("\\textbf{",mean_fd,"}"), mean_fd
  mutate(Stabilization = paste0(Stabilization,"\\quad{\\footnotesize(",mean_sd-2*sem_sd, "--", mean_sd+
         `Fitness Difference` = paste0(`Fitness Difference`," {(\\footnotesize",mean_fd-2*sem_fd, "--",
  select(`Species pair`,
         Code = pair, Stabilization,
         `Fitness Difference`,
         `Net effect of PSF` = outcome2) %>%
  # arrange(`Species pair`) %>%
  mutate_all(kableExtra::linebreak, align = "c")

tbl_to_print <- kableExtra::kable(tbl_to_print,  booktabs = T, escape = F, align = "lcccc", format = "la
  kableExtra::kable_styling(font_size = 10.5)  %>%
  kableExtra::column_spec(1, width = "1.35in") %>%
  kableExtra::column_spec(3, width = ".85in") %>%
  kableExtra::column_spec(4, width = ".85in") %>%
  kableExtra::pack_rows("Plant-soil feedbacks tend to promote coexistence (stabilization > fitness diff
                        latex_gap_space = "1em") %>%
  kableExtra::pack_rows("Plant-soil feedbacks tend to promote exclusion (fitness difference > stabiliza
                        indent = F, latex_gap_space = "1em") %>%
  kableExtra::pack_rows("Strong evidence that plant-soil feedbacks promote exclusion\n(lower bound fitn
                        indent = F, latex_gap_space = "1em") %>%
  kableExtra::row_spec(c(1,4,6,8,10,12,14), extra_latex_after = "\\rowcolor{gray!10}")

if(write_objects){
  saveRDS(tbl_to_print, "manuscript/figures/outcomes_table.Rds")
}
```

**Elements of the supplementary materials.**

First, we analyze data from the Variation experiment for Supplement 2:

```
# Variation experiment -----
var_expt <- read_csv("../../data/variation-harvest.csv")
```

```
## Parsed with column specification:
## cols(
##   pot_id = col_double(),
##   pot_tag = col_character(),
##   experiment_id = col_character(),
##   source_soil = col_character(),
##   focal_species = col_character(),
##   replicate = col_character(),
##   abg_dry_g = col_double()
## )
```

```r
# Two plants died and have NA in abg_dry_g, let's eliminate these
num_NA_pots_var <- var_expt %>% filter((is.na(abg_dry_g))) %>% nrow

var_expt <- var_expt %>% filter(!(is.na(abg_dry_g)))

# Make a dataset of median, lqr, uqr, and outlier
# as we had done for the main experiment
variation_expt_data_sum <- var_expt %>%
  group_by(source_soil, focal_species) %>%
  summarize(median_bm = median(abg_dry_g),
            lqr = quantile(abg_dry_g, 0.25),
            uqr = quantile(abg_dry_g, 0.75),
            min_val = min(abg_dry_g, na.rm = T),
            max_val = max(abg_dry_g, na.rm = T),
            out_low = ifelse(min_val < lqr - 1.5*(uqr-lqr), min_val, NA),
            out_upp = ifelse(max_val > uqr + 1.5*(uqr-lqr), max_val, NA),
            n = n())

# I want to also extract the rows for PL_PLER and PL_FEMI from the
# dataframe used for the earlier dotplot.
# Additionally I want to show the averages from the variation experiment
variation_expt_data_splevel <- var_expt %>%
  group_by(focal_species) %>%
  summarize(median_bm = median(abg_dry_g),
            lqr = quantile(abg_dry_g, 0.25),
            uqr = quantile(abg_dry_g, 0.75),
            min_val = min(abg_dry_g, na.rm = T),
            max_val = max(abg_dry_g, na.rm = T),
            out_low = ifelse(min_val < lqr - 1.5*(uqr-lqr), min_val, NA),
            out_upp = ifelse(max_val > uqr + 1.5*(uqr-lqr), max_val, NA),
            n = n()) %>%
  mutate(source_soil = "PL_separate_averaged") %>%
  select(source_soil, everything())

var_expt_for_cleveland <- bind_rows(variation_expt_data_sum,
                                    # The rows for PLER and FEMI in PL soil in the main experiment
                                    # variation_expt_data_splevel,
                                    biomass_for_cleveland %>%
                                      filter(focal_species %in% c("PLER", "FEMI"),
                                             source_soil == "PL"),
                                    .id = "which_df") %>%
  arrange(focal_species) %>% ungroup %>%
  mutate(plot_y = c(seq(0.8,1.2, length.out = 6),
                    seq(1.8, 2.2, length.out = 6)))

# Make the "Cleveland plot" for the variation experiment -----
(var_expt_cleveland  <- ggplot(var_expt_for_cleveland,
                               aes(x = median_bm, y = plot_y,
                                   shape = which_df, fill = which_df)) +
   scale_x_log10() +
   # add median value
   geom_point(size = 4, stroke = .9) +
```

```
scale_shape_manual(name = "Inoculum source",
                   values = c(21, 24), #, 22),
                   labels = c("Single PLER\nmonoculture",
                              "Homogenized across\nPLER monocultures")) +
scale_fill_manual(name = "Inoculum source",
                  values = c("#56B4E9", "#F0E442"),#, "#CC79A7"),
                  labels = c("Single PLER\nmonoculture",
                             "Homogenized across\nPLER monocultures")) +
# add dashed lines connecting outlier to median (if outlier exists)
geom_errorbarh(aes(xmin = out_low, xmax = lqr),
               height = 0, linetype = 3, size = .25) +
geom_errorbarh(aes(xmin = uqr, xmax = out_upp),
               height = 0, linetype = 3, size = .25) +
# add outlying points (if they exist)
geom_point(aes(y = plot_y, x = out_low), shape = 21, fill = "grey50", size = .5) +
geom_point(aes(y = plot_y, x = out_upp), shape = 21, fill = "grey50", size = .5) +
# add solid line connecting median to lower and upper quantile
geom_errorbarh(aes(xmin = lqr, xmax = uqr), height = .01) +
theme_gsk() +
theme(legend.justification=c(0,0), legend.position=c(.65,.40),
      # legend.position = "right",
      legend.background = element_rect(colour = "grey50"),
      legend.text = element_text(size = 7),
      legend.title = element_text(size = 8)
) +
xlab("Aboveground biomass (g)") +
ylab("Focal species") +
scale_y_continuous(breaks = c(1,2),
                   labels = c("FEMI", "PLER")) +

NULL)
```

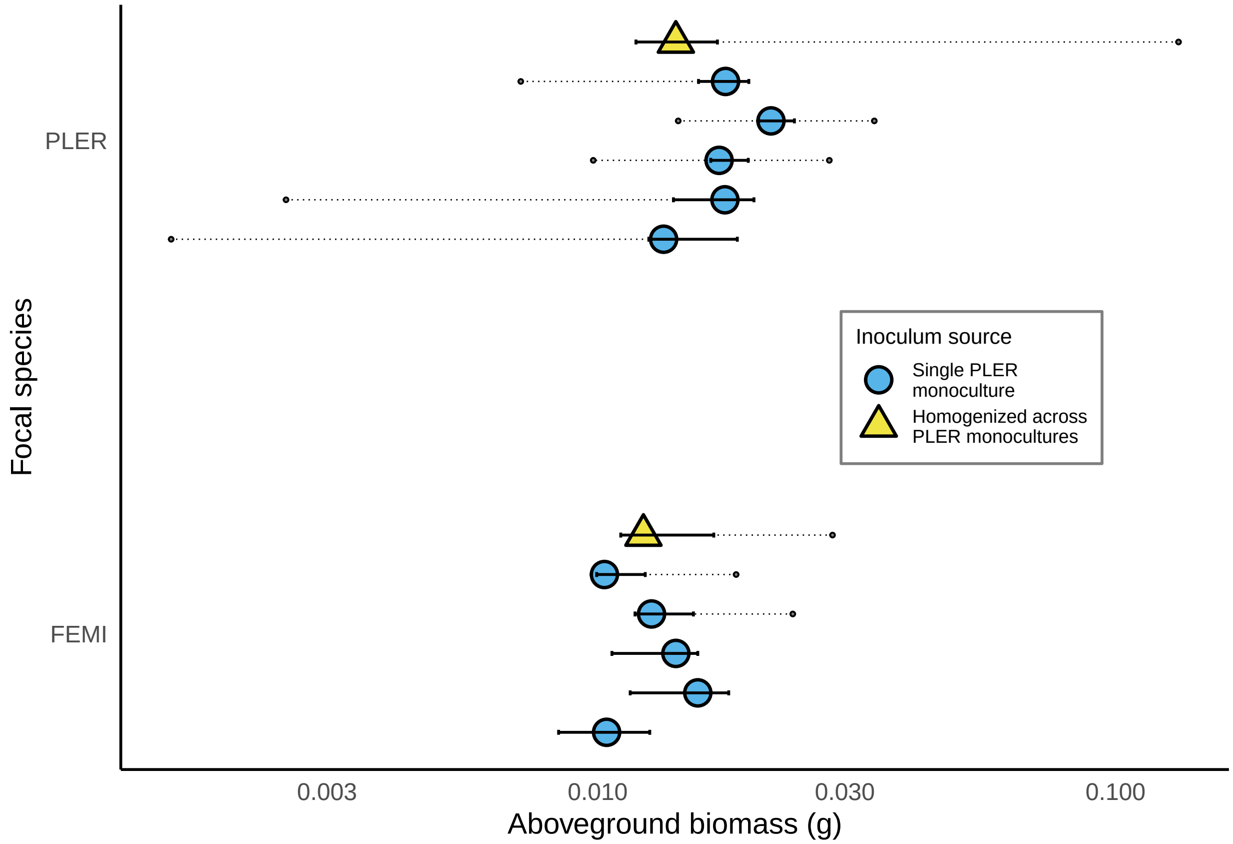

```
if(write_objects) {
  saveRDS(var_expt_cleveland, file = "../../figures/var_expt_cleveland.Rds")
}
# Compare variances across groups in Var Expt ------
# The goal here is to ask whether variance is higher in
# the 50 plants from the variation experiment vs. the 10 PL_PLER
# or PL_FEMI plants.

var_and_main <- bind_rows(var_expt %>% rename(number = pot_id) %>% select(experiment_id, everything()),
                          biomass %>% rename(pot_tag = pot_id) %>%
                            filter(focal_species %in% c("PLER", "FEMI"),
                                   source_soil == "PL") %>%
                            select(-pointcol) %>% mutate(experiment_id = "main") %>%
                            select(experiment_id, everything())) %>%
  mutate(experiment_id = as.factor(experiment_id))
var_and_main_pl <- var_and_main %>% filter(focal_species == "PLER")

summary(aov(log(abg_dry_g)~source_soil, data = var_and_main %>% filter(focal_species == "PLER")))
```

```
##              Df Sum Sq Mean Sq F value Pr(>F)
## source_soil   5   1.75   0.350    1.16   0.34
## Residuals    55  16.56   0.301
```

```
pl_var_leveneP <- car::leveneTest(abg_dry_g~source_soil, data = var_and_main %>% filter(focal_species =
pl_var_leveneP
```

```
## Levene's Test for Homogeneity of Variance (center = median)
##       Df F value Pr(>F)
```

```
## group  5    0.82   0.54
##       55
```

```
summary(aov(log(abg_dry_g)~source_soil, data = var_and_main %>% filter(focal_species == "FEMI")))
```

```
##             Df Sum Sq Mean Sq F value Pr(>F)
## source_soil  5   0.73  0.1464    1.78   0.13
## Residuals   52   4.28  0.0823
```

```
fe_var_leveneP <- car::leveneTest(abg_dry_g~source_soil, data = var_and_main %>% filter(focal_species =
fe_var_leveneP
```

```
## Levene's Test for Homogeneity of Variance (center = median)
##       Df F value Pr(>F)
## group  5    0.44   0.82
##       52
```

Finally, comparing the inference of coexistence/exclusion based on our approach (Eqn. 3 of the main text) vs. the inference based on the feasibility analysis of Bever et al. (1997).

```
# Comparing ND/FD decomp to feasibility analysis ---------
biomass <- read_csv("../../data/phase2-harvest.csv")
```

```
## Parsed with column specification:
## cols(
##   number = col_double(),
##   pot_id = col_character(),
##   source_soil = col_character(),
##   focal_species = col_character(),
##   replicate = col_character(),
##   abg_dry_g = col_double()
## )
```

```
# Get rid of "empty" rows
biomass <- biomass %>% arrange(focal_species, source_soil, replicate) %>%
  filter(!(grepl("Empty", biomass$source_soil)))
biomass$focal_species_2let <- str_extract(biomass$focal_species, "..")
biomass$log_agb <- log(biomass$abg_dry_g)
combinations <- combn(unique(biomass$focal_species_2let),2)
biomass_arranged <- tibble(pair = NA,
                           rep  = NA,
                           B1A = NA, B1B = NA, B1F = NA, B2A = NA, B2B = NA, B2F = NA)

for(current_combination in 1:ncol(combinations)) {
  pair <- combinations[,current_combination]
  sp1 <- pair[1]; sp2 <- pair[2]

  # subset the biomass data file
  biomass_sub <-
    biomass %>%
    filter(source_soil %in% c(pair, "abfld") & focal_species_2let %in% pair)

  bm1_1 <- biomass_sub %>% filter(source_soil == pair[1], focal_species_2let == pair[1]) %>%
    select(log_agb) %>% unlist
  bm1_2 <- biomass_sub %>% filter(source_soil == pair[2], focal_species_2let == pair[1]) %>%
    select(log_agb) %>% unlist
```

```
  bm1_f <- biomass_sub %>% filter(source_soil == "abfld", focal_species_2let == pair[1]) %>%
    select(log_agb) %>% unlist

  bm2_1 <- biomass_sub %>% filter(source_soil == pair[1], focal_species_2let == pair[2]) %>%
    select(log_agb) %>% unlist
  bm2_2 <- biomass_sub %>% filter(source_soil == pair[2], focal_species_2let == pair[2]) %>%
    select(log_agb) %>% unlist
  bm2_f <- biomass_sub %>% filter(source_soil == "abfld", focal_species_2let == pair[2]) %>%
    select(log_agb) %>% unlist

  new_df <- tibble(pair = paste0(pair[1], "_", pair[2]),
                   rep  = unique(biomass_sub$replicate),
                   B1A = bm1_1, B1B = bm1_2, B1F = bm1_f,
                   B2A = bm2_1, B2B = bm2_2, B2F = bm2_f)
  biomass_arranged <- bind_rows(biomass_arranged, new_df)
}
biomass_arranged <- biomass_arranged %>% filter(!is.na(pair))
predict_each_rep <- biomass_arranged %>%
  mutate(IS = (B1A-B1F) - (B1B-B1F) - (B2A-B2F) + (B2B-B2F)) %>%
  mutate(stabilization = -0.5*IS) %>%
  mutate(fitdiff = 0.5*((B1A-B1F) + (B1B-B1F) - (B2A-B2F) - (B2B-B2F))) %>%
  mutate(p1_num = ((B2B-B2F) - (B1B-B1F))) %>%
  mutate(p2_num = ((B1A-B1F) - (B2A-B2F))) %>%
  mutate(check = (p1_num+p2_num - IS < .001)) %>%
  mutate(p1 = p1_num/IS, p2 = p2_num/IS) %>%
  mutate(coex_stfd = ifelse(abs(fitdiff) < stabilization, "coex", "exclude")) %>%
  mutate(coex_IS = ifelse(IS < 0 & p1*p2 > 0, "coex", "exclude"))
# predict_each_rep %>% View
which(predict_each_rep$coex_stfd != predict_each_rep$coex_IS)
```

```
## integer(0)
```

```
table_s1.1 <- predict_each_rep %>%
  mutate(m1A = B1A-B1F, m1B = B1B-B1F,
         m2A = B2A-B2F, m2B = B2B-B2F,
         pair = str_replace(pair, "_", "\\\\_")) %>%
  mutate_if(is.numeric, round, 3) %>%
  select( rep, pair,
         `$m_{1A}$` = m1A, `$m_{1B}$` = m1B,
         `$m_{2A}$` = m2A, `$m_{2B}$` = m2B,
         `$I_S$` = IS,
         `$\\hat{p_1}$` = p1, `$\\hat{p_2}$` =p2,
         `outcome \n (feasibility)` = coex_IS,
          stabilization,
         `fitness difference` = fitdiff, `outcome (IGR)` = coex_stfd) %>%
  filter(!is.na(`outcome (IGR)`)) %>%
  mutate(rep = str_remove(rep, "R"),
         rep = as.numeric(rep)) %>%
  arrange(pair, rep) %>%
  mutate_all(kableExtra::linebreak)  %>%
  kableExtra::kable(booktabs = T, longtable = T,
                    linesep = c(rep("",9),  "\\addlinespace\\addlinespace",
                                rep("",9),  "\\addlinespace\\addlinespace",
                                rep("",9),  "\\addlinespace\\addlinespace",
```

```
                                        rep("",7),  "\\addlinespace\\addlinespace",
                                        rep("",7),  "\\addlinespace\\addlinespace",
                                        rep("",9),  "\\addlinespace\\addlinespace",
                                        rep("",9),  "\\addlinespace\\addlinespace",
                                        rep("",7),  "\\addlinespace\\addlinespace",
                                        rep("",7),  "\\addlinespace\\addlinespace",
                                        rep("",7),  "\\addlinespace\\addlinespace",
                                        rep("",7),  "\\addlinespace\\addlinespace",
                                        rep("",7),  "\\addlinespace\\addlinespace",
                                        rep("",7),  "\\addlinespace\\addlinespace",
                                        rep("",7),  "\\addlinespace\\addlinespace",
                                        rep("",5)),
                         escape = F,format = "latex", align = "cccccc|cccc|ccc") %>%
  kableExtra::column_spec(c(10,12,13), width = ".1in") %>%
  kableExtra::kable_styling(font_size = 10)  %>%
  kableExtra::row_spec(c(10:19, 30:37, 46:55, 66:73, 82:89, 98:105, 114:121), extra_latex_after = "\\ro

if(write_objects){
  saveRDS(table_s1.1, "../../figures/table_s1.1.Rds")
}
```
